# Elucidation of remdesivir cytotoxicity pathways through genome-wide CRISPR-Cas9 screening and transcriptomics

**DOI:** 10.1101/2020.08.27.270819

**Authors:** Ersin Akinci, Minsun Cha, Lin Lin, Grace Yeo, Marisa C. Hamilton, Callie J. Donahue, Heysol C. Bermudez-Cabrera, Larissa C. Zanetti, Maggie Chen, Sammy A. Barkal, Benyapa Khowpinitchai, Nam Chu, Minja Velimirovic, Rikita Jodhani, James D. Fife, Miha Sovrovic, Philip A. Cole, Robert A. Davey, Christopher A. Cassa, Richard I. Sherwood

**Author notes:** Corresponding author: R.I.S., Correspondence. These authors contributed equally to this work (Ersin Akinci, Minsun Cha, Lin Lin, Grace Yeo, Marisa C. Hamilton). These authors contributed equally to this work (Callie J. Donahue, Heysol C. Bermudez-Cabrera, Larissa C. Zanetti, Maggie Chen, Sammy A. Barkal, Benyapa Khowpinitchai).

## Abstract

The adenosine analogue remdesivir has emerged as a frontline antiviral treatment for SARS-CoV-2, with preliminary evidence that it reduces the duration and severity of illness^1^. Prior clinical studies have identified adverse events^1,2^, and remdesivir has been shown to inhibit mitochondrial RNA polymerase in biochemical experiments^7^, yet little is known about the specific genetic pathways involved in cellular remdesivir metabolism and cytotoxicity. Through genome-wide CRISPR-Cas9 screening and RNA sequencing, we show that remdesivir treatment leads to a repression of mitochondrial respiratory activity, and we identify five genes whose loss significantly reduces remdesivir cytotoxicity. In particular, we show that loss of the mitochondrial nucleoside transporter *SLC29A3* mitigates remdesivir toxicity without a commensurate decrease in SARS-CoV-2 antiviral potency and that the mitochondrial adenylate kinase *AK2* is a remdesivir kinase required for remdesivir efficacy and toxicity. This work elucidates the cellular mechanisms of remdesivir metabolism and provides a candidate gene target to reduce remdesivir cytotoxicity.

---

Remdesivir is a phosphoramidate prodrug which is converted within cells to an adenosine triphosphate analogue^2^. This active remdesivir metabolite is incorporated by viral RNA-dependent RNA polymerase (RdRp) enzymes into nascent viral RNA chains and results in premature transcript termination^3^. While developed to treat Ebola virus^3^, remdesivir has been shown to impede replication of coronaviruses including SARS-CoV and MERS-CoV^4,5^ and notably SARS-CoV-2^6–8^ *in vitro* and in mouse and macaque models.

While clinical data suggests remdesivir is generally safe for human use, the causes and mechanisms of toxicity are not fully known. These may depend on patient-specific conditions or genetic background, and may increase with higher dosing or treatment earlier in disease progression as has been shown for other antiviral drugs such as oseltamivir (Tamiflu)^1^,^5^,^7^,^9–11^. One possible mode of toxicity induced by nucleoside analogues is mitochondrial toxicity, as mitochondrial polymerases lack the selectivity of mammalian polymerases to exclude nucleoside analogues. HIV antiviral nucleotide analogues and Hepatitis C virus (HCV) nucleoside analogs induce mitochondrial toxicity with varying levels of severity^12,13^, and remdesivir has been shown to inhibit mitochondrial RNA polymerase in biochemical experiments^14^, albeit at 100-fold lower rates than RdRp.

In order to better understand the mechanisms by which remdesivir affects host cells and induces cytotoxicity, we have performed RNA-sequencing (RNA-seq) and genomewide CRISPR-Cas9 screening on liver and intestinal cell lines treated with remdesivir. RNA-seq has been employed previously to characterize drug mechanisms, which include cytotoxicity^15^,^16^. CRISPR-Cas9 screening has been used to identify genetic pathways driving cancer drug resistance^17^. To our knowledge, neither approach has been used to characterize the effects of remdesivir.

To begin addressing which genes and pathways underlie remdesivir toxicity, we performed RNA-seq in two human intestinal cell lines (HT29 and HCT116) and two human liver cell lines (HepG2 and PLC/PRF/5). We use these cell lines because they are derived from organs that are often most affected by drug toxicity. We performed RNA-seq on all four cell lines harvested after 8 and 24 hours of remdesivir treatment using a maximal dose for each cell line which does not induce apoptosis within a 24 hour measurement time (Supplementary Table 1). As controls, we performed similar experiments on DMSO-treated cells and cells treated with the SARS-CoV-2 antiviral drug hydroxychloroquine (HCQ)^18^.

We hypothesized that HCQ would induce distinct host gene expression response due to its orthogonal mechanism of impairing endosome and lysosome trafficking^19^.

As expected, RNA-seq gene expression clusters most strongly by cell line and next by drug exposure (Fig. 1a, Supplementary Fig. 1). There is clear evidence of differential expression without strong pro-apoptotic gene expression signatures, validating our dosing and duration (Supplementary Fig. 1). To investigate which genes and pathways change expression as a result of remdesivir treatment, we performed gene set enrichment analysis (GSEA)^20^. We identify a set of highly statistically significant gene ontology (GO) pathways affected by remdesivir in each of the four cell lines. There is concordance in the pathways regulated by remdesivir across the four cell lines that differ from pathways induced by HCQ treatment (Fig. 1b, Supplementary Fig. 1), suggesting that each drug has a signature impact on cellular physiology.

**Figure 1.**
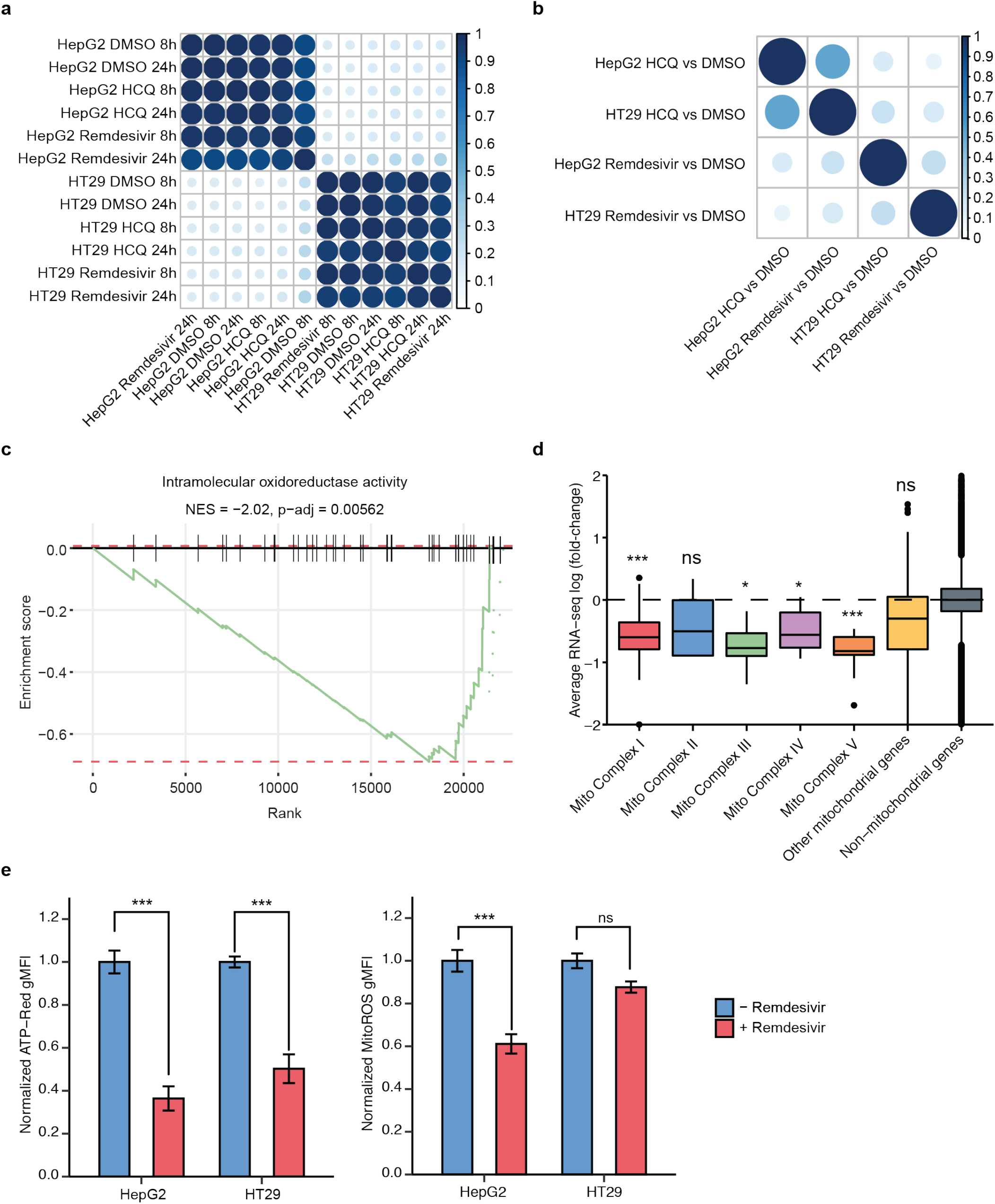
RNA-seq upon remdesivir treatment reveals decreased expression of genes involved in mitochondrial respiration. a) RNA-seq gene expression correlation matrix among HT29 and HepG2 cell lines treated with remdesivir, HCQ, or DMSO (control) for 8 and 24 hours. b) GSEA correlation matrix of HT29 and HepG2 cell lines treated with remdesivir or HCQ relative to DMSO (control). c) GSEA enrichment plot of intramolecular oxidoreductase activity in HT29 for remdesivir treatment as compared to DMSO control. d) Average mitochondrial RNA-seq log2(fold-change) of HepG2 cells after 24 hour treatment with remdesivir as compared to DMSO control. e) Normalized ATP-red and mitoROS geometric mean of fluorescence intensity (gMFI) for untreated (blue) or remdesivir-treated (red) HepG2 and HT29 cells.

The main drug-specific gene expression response of remdesivir is a significant decrease in mitochondrial respiratory gene expression (Fig. 1c-d, Supplementary Fig. 1, Supplementary Table 2). Remdesivir strongly represses nuclear genes that comprise mitochondrial respiratory complexes 1, 3, 4, and 5 as well as other genes encoding mitochondrially localized proteins while not significantly impacting expression of genes encoded by the mitochondrial genome (Fig. 1d, Supplementary Fig. 1). To determine the functional impact of remdesivir mitochondrial gene repression, we performed flow cytometric measurement of mitochondrial redox state and ATP production using the fluorescent dyes MitoROS 520 and ATP-Red^21^,^22^, finding a significant decrease in ATP production upon remdesivir treatment in HepG2 and HT29 cells and a significant decrease in mitochondrial oxidation in HepG2 cells (Fig. 1e-f). Thus, we conclude that remdesivir decreases mitochondrial function, at least partially through decreasing the expression of genes involved in mitochondrial respiration.

The main cell line-consistent remdesivir-upregulated pathways are the RNA polymerase and nutrient stress response pathways driven by ATF3 and ATF4, which are upregulated in all cell lines (Supplementary Table 2)^23^ and which should dampen mTOR signaling^24^. These pathways are also up-regulated by HCQ and are a common cellular response to stress, including mitochondrial stress^25^. In contrast to remdesivir, HCQ robustly upregulates cholesterol metabolism, autophagy, and starvation pathways and decreases rRNA and tRNA production across all cell lines (Supplementary Table 2), consistent with the known HCQ mechanism of impairment of endosome to lysosome processing^19^,^26^ and suggesting that RNA-seq analysis highlights specific drug-response pathways. We show that HCQ impairs cellular LDL uptake and induces an endosomal LDL retention phenotype (Supplementary Fig. 2), confirming this consistent phenotypic outcome in the HCQ RNA-seq.

Given that remdesivir decreases the expression of genes involved in mitochondrial respiration, we asked whether drugs that affect mitochondrial oxidative phosphorylation impact remdesivir toxicity. We tested a set of five drugs previously reported to impact mitochondrial function, finding that remdesivir cytotoxicity in HT29 and HepG2 cells is mitigated by syrosingopine, exacerbated by ursodeoxycholic acid, and unaffected by trigonelline, dimethyl fumarate, and metformin (Supplementary Fig. 3). Ursodeoxycholic acid has been shown to improve mitochondrial function^27^, and syrosingopine, a lactate transporter inhibitor, has been shown to decrease mitochondrial function^28,29^. These results suggest that further suppressing mitochondrial activity may mitigate remdesivir toxicity, although the mitochondrial suppressor metformin did not replicate this activity, suggesting that there are nuances to how altering mitochondrial activity impacts remdesivir cytotoxicity.

In order to directly identify genes involved in remdesivir metabolism and cytotoxicity, we developed a drug toxicity CRISPR-Cas9 screening paradigm. A pool of HT29 cells was treated with the Brunello lentiviral genome-wide CRISPR-Cas9 gene knockout library^30^, which targets 19,114 genes each with four guide RNAs (gRNAs), and after selection for cells with lentiviral integration, cells were treated for 14 days either with remdesivir or with DMSO or HCQ as controls. We used concentrations of remdesivir and HCQ that yield 50% as many surviving cells every five days (Supplementary Fig. 4) to detect genes whose knockout exacerbates drug toxicity (relatively depleted in remdesivir-treated samples) or mitigates toxicity (enriched) (Fig. 2a). To address whether this screening paradigm indeed measures remdesivir-dependent toxicity, we treated HT29 cells with remdesivir in the presence of a 100-fold molar excess of adenosine, which should decrease the relative usage of remdesivir by mitochondrial RNA polymerase, finding that remdesivir toxicity is completely attenuated (Fig. 2b).

**Figure 2.**
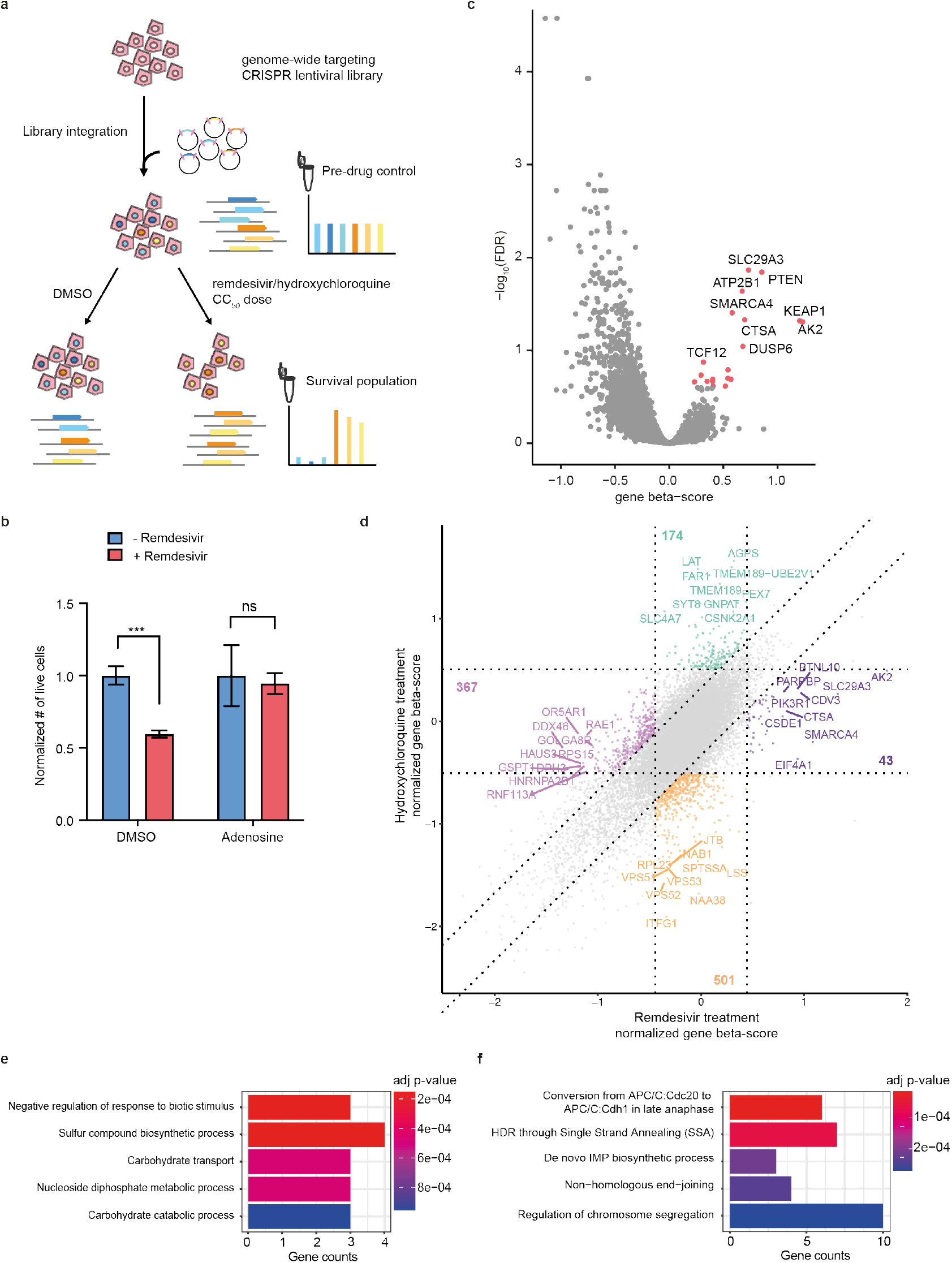
A drug toxicity CRISPR-Cas9 screen identifies candidate genes and pathways involved in the mitigation and exacerbation of remdesivir cytotoxicity. a) Schematic of pooled CRISPR screen. Cells were transduced with Brunello lentiviral library and treated either with DMSO or for 14 days with doses of remdesivir or HCQ titrated to yield 50% surviving cells every five days of treatment. Genomic DNA was collected before and after drug for sequencing of gRNA abundance. b) Effect on remdesivir toxicity of 100-fold molar excess of adenosine in HT29 cells. Error bars are SE. c) Volcano plot showing beta score calculated for the 4 gRNAs targeting each gene vs. the −log_10_(FDR) of remdesivir treatment vs. DMSO treatment across 6 screening replicates. d) Square plot showing normalized gene beta score normalized to pre-drug control for remdesivir vs. HCQ treatment. e-f) Top 5 most significant GO terms enriched (e) and depleted (f) after remdesivir treatment vs. DMSO treatment.

We performed six biological replicate screens in HT29 cells with both drugs and used the MAGeCK-MLE pipeline^31^,^32^ to identify genes and pathways mediating antiviral drug toxicity (Supplementary Table 3). We collected an average of 5.5∗10^6 NGS reads per library, obtaining high replicate consistency in gRNA representation (r = 0.60-0.77, Supplementary Fig. 4). We observed a significant depletion of gRNAs targeting cell essential genes after two weeks of drug or DMSO treatment, confirming robust CRISPR-Cas9 activity (Supplementary Fig. 4). To identify gRNAs specifically enriched or depleted by drug treatment, we calculated the MAGeCK beta-score per gene compared to DMSO control to identify significantly enriched and depleted genes (Fig. 2c, Supplementary Fig. 4), then compared the remdesivir and HCQ screens to identify sets of 43-501 genes specifically affected by each drug (Fig. 2d, Supplementary Fig. 4).

To discover pathways regulating remdesivir metabolism and toxicity, we performed GO analysis on these genes whose knockout specifically alters growth in the presence of remdesivir or HCQ. We found that genes involved in carbohydrate transport and catabolism as well as nucleoside diphosphate metabolism are enriched after remdesivir treatment (Fig. 2e).

These pathways match known steps in the enzymatic activation of phosphoramidate prodrugs such as remdesivir^33^. Genes involved in the synthesis of the purine precursor inosine monophosphate (IMP), including ATIC, GART, and PFAS, emerge as among the most depleted after remdesivir treatment (Fig. 2f), likely by restricting cellular concentrations of adenosine and thus increasing relative abundance of remdesivir. DNA repair-related pathways are also depleted after remdesivir treatment.

On the gene level, we performed individual validation of gRNAs targeting six genes whose loss significantly mitigated remdesivir toxicity in the pooled screen, confirming significantly increased CC_50_ in five out of six (Fig. 3a, Supplementary Fig. 5). We were unable to maintain knockout cells for the sixth gene, EIF4A1, because of excessive cell death. The five confirmed genes elucidate mechanisms of remdesivir activation and toxicity. Among these genes is cathepsin A (CTSA, 11.3-fold CC_50_ increase in HT29), an esterase known to be involved in intracellular activation of phosphoramidate nucleoside prodrugs^29^, providing genetic confirmation that CTSA is the primary enzyme in the initial intracellular remdesivir activation step. Additionally, gRNAs targeting Kelch-like ECH-associated protein 1 (KEAP1, 3.2-fold CC_50_ increase in HT29), whose knockout should stabilize and activate the NRF2 transcription factor, were enriched. Activated NRF2 is established to increase a set of antioxidant response element genes involved in stress response, mitochondrial function, and purine synthesis^34^, suggesting that NRF2 activation may protect against remdesivir toxicity. The plasma membrane calcium transporter ATP2B1 (5.7-fold CC_50_ increase in HT29) was also required for remdesivir toxicity through an unknown mechanism.

**Figure 3.**
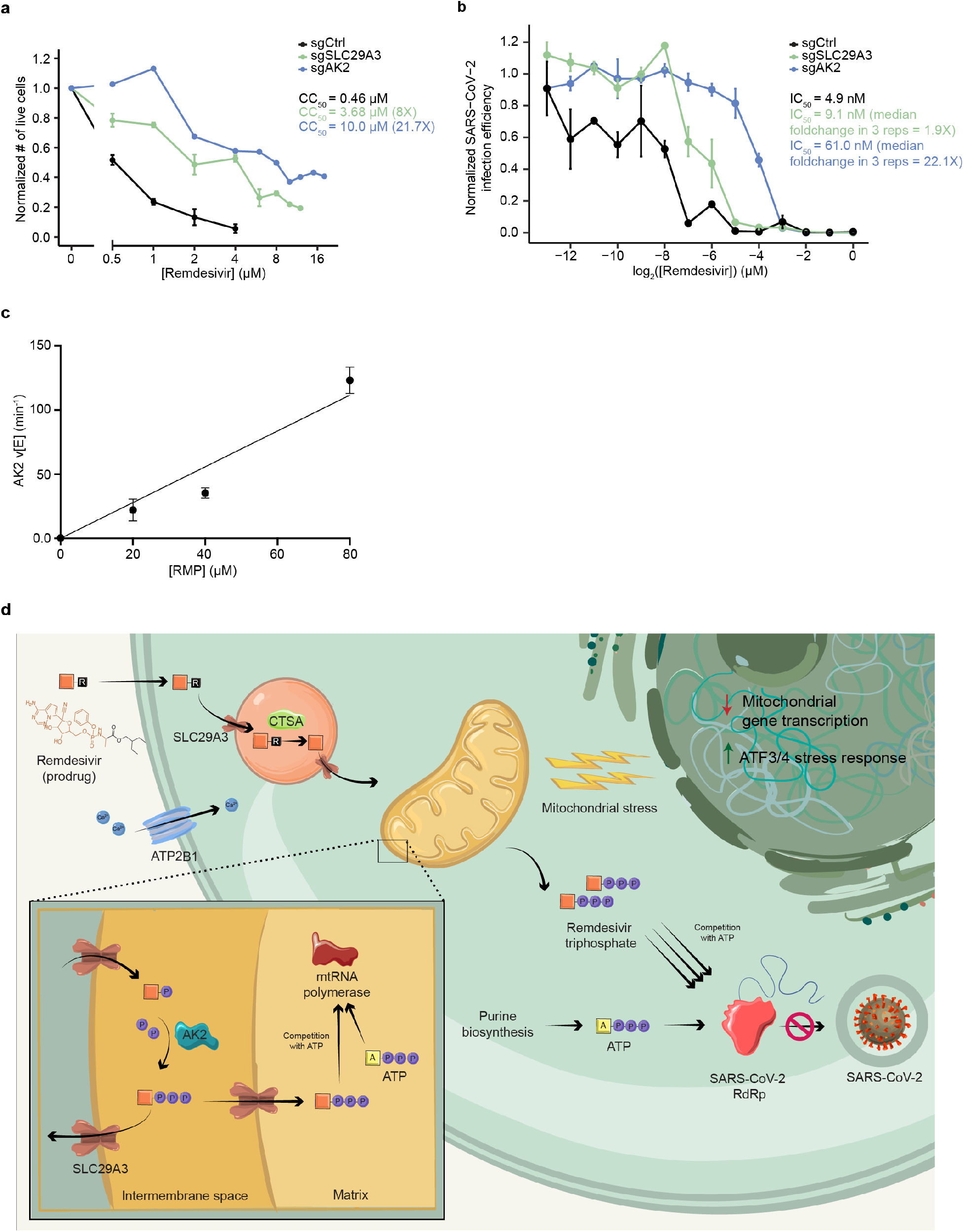
*SLC29A3* knockout mitigates remdesivir toxicity without commensurate loss of SARS-CoV-2 antiviral potency, and AK2 is a remdesivir kinase required for antiviral potency and cytotoxicity. a) Effect of remdesivir on average normalized number of live cells and CC_50_ in Huh7 sgCtrl and KO cell lines. b) Effect of remdesivir on average normalized SARS-CoV-2 infection efficiency and IC_50_ in sgCtrl and KO cell lines. c) Steady-state kinetic plot for AK2 catalytic rate (v/[E]) versus [RMP] (0-80 *μ* M). Reactions were carried out with 40 nM AK2 at 30° C for 10 minutes. d) Schematic of genes and pathways involved in the cellular metabolism and toxicity of remdesivir. The remdesivir prodrug is metabolized sequentially by Cathepsin A in the lysosome and AK2 in the mitochondrial intermembrane space into its active remdesivir-triphosphate form, which restricts SARS-CoV-2 RdRp in competition with ATP. SLC29A3 may modulate remdesivir-triphosphate entry into the mitochondrial matrix where its inhibition of mitochondrial RNA polymerase leads to toxicity. Remdesivir treatment leads to repressed nuclear transcription of mitochondrial genes, especially those involved in respiration, as well as leading to an ATF3/4 stress response. The plasma membrane transporter ATP2B1 is also involved in remdesivir cytotoxicity through an unknown mechanism.

Intriguingly, gRNAs targeting two mitochondrially localized adenosine metabolic genes, the reversible adenylate kinase AK2^35,36^ and the bidirectional nucleoside transporter SLC29A3^37^, are highly enriched. Notably, AK2 has been shown to be involved in activation of the antiviral drugs tenofovir and PMEA^38,39^, and *SLC29A3* knockdown has been shown to reduce mitochondrial transport of other nucleotide analogue drugs^17^, suggesting that both of these genes are common regulators of antiviral nucleotide drug metabolism. We found that the CC_50_ for remdesivir is dramatically increased by knockout of AK2 (71.4, 58.9, 21.7, 9.6, and 5.9-fold) and SLC29A3 (11.3, 9.8, 8.0, 5.7, and 10.9-fold) in HepG2, HT29, Huh7, OV-90, and HEK293-ACE2 cells respectively (Fig. 3a, Supplementary Fig. 5), suggesting that these genes are common cellular mediators of remdesivir toxicity. In contrast to their reported impacts on metabolism of other antiviral nucleotide analogue drugs^37–39^, knockout of *AK2* and SLC29A3 did not significantly impact the cytotoxicity of the cytosine mimetic SARS-CoV-2 antiviral EIDD-1931^40^ in HT29 cells (Supplementary Fig. 5), which may be because EIDD-1931 causes cytotoxicity through mechanisms other than mitochondrial polymerase inhibition^41^.

Additional genes whose knockout mitigates remdesivir toxicity include EIF4A1, TSC1, TSC2, DUSP4, and DUSP6, whose knockouts should all activate mTOR signaling^42,43^, suggesting that activating mTOR signaling may counteract the ATF3- and ATF4-driven stress response observed in the RNA-seq data. Overall, remdesivir cytotoxicity can be mitigated by modulating conversion of prodrug to active drug, mitochondrial nucleoside metabolism and transport, mitochondrial function, and mTOR signaling, and cytotoxicity is exacerbated by blocking synthesis of purine nucleotides that would compete with remdesivir.

In contrast, an orthogonal and highly biologically coherent set of genes impacts the cytotoxicity of HCQ treatment. Nearly all of the genes most enriched genes after HCQ treatment are involved in plasmalogen synthesis in the peroxisome (core plasmalogen synthesis genes AGPS1, FAR1, and TMEM189^44^ are the three most enriched genes), peroxisome maintenance (e.g. PEX5, PEX6, and PEX7^45^), or vesicle trafficking (e.g. SYT8, RALGAPB, and CHD1) (Supplementary Fig. 6), which is supported by GO analysis (Supplementary Fig. 6). Genes depleted by HCQ are highly enriched for those involved with endocytosis, cholesterol metabolism, and vesicle trafficking (Supplementary Fig. 6). These results are consistent with the RNA-seq data indicating that HCQ inhibits cholesterol metabolism, suggesting that further impairing cholesterol production is synthetic lethal to HCQ-treated cells. They are also concordant with the known role of HCQ in disrupting endosome to lysosome processing^19,26^. We confirmed 5/7 gene knockouts individually, showing changes in survival upon HCQ treatment consistent with screening results (Supplementary Fig. 6), and we found that knockout of the sterol biosynthesis gene *SQLE* reversed the HCQ resistance induced by loss of the plasmalogen synthesis genes *AGPS* and *FAR*1 (Supplementary Fig. 6), supporting the idea that plasmalogens increase HCQ cytotoxicity through inhibiting cholesterol production^44^.

To address whether loss of *AK2* or *SLC29A3* impacts the potency of remdesivir as a SARS-CoV-2 antiviral agent, we tested remdesivir response in a SARS-CoV-2 virus challenge assay in the presence of individual gene knockout in Huh7 cells. Control or knockout cells were treated with a range of remdesivir concentrations and then infected with SARS-CoV-2 viral supernatant for 48 hours before fixation and RNA fluorescence *in situ* hybridization detection of SARS-CoV-2 RNA (Supplementary Fig. 7). We found in three independent replicate virus challenge experiments that the SARS-CoV-2 IC_50_ for remdesivir in Huh7 is increased only slightly by *SLC29A3* knockout as compared with control gRNA (median 1.9-fold), while AK2 knockout induces a robust (median 22.1-fold) increase in the remdesivir IC_50_ (Fig. 3b, Supplementary Fig. 7). The increase in IC_50_ induced by *AK2* knockout in Huh7 is highly similar to the increase in CC_50_ dose (21.7-fold), suggesting that loss of *AK2* decreases cytotoxicity through decreasing the intracellular availability of active remdesivir-triphosphate. However, SLC29A3 loss induces a robust decrease in cytotoxicity (8-fold) without a commensurate decrease in antiviral potency.

Given the requirement of *AK2* in SARS-CoV-2 antiviral efficacy and its role in phosphorylating other nucleoside analog drugs^38^,^39^, we asked whether AK2 is a remdesivir kinase. After the remdesivir prodrug is uncaged by CTSA and cellular amidases, it forms remdesivir monophosphate (RMP)^33,46,47^. As RMP requires phosphorylation to actively inhibit RdRp or mitochondrial RNA polymerase, we examined the possibility that AK2 can convert RMP to the diphosphate (RDP) form using a spectrophotometric pyruvate kinase/lactate dehydrogenase coupled assay. This enzyme assay connects any ADP formation that occurs upon phosphorylation of RMP to the oxidation of NADH, which can be followed by UV absorbance change at 340 nm (Supplementary Fig. 8)^48–50^. Using this assay, we demonstrated that recombinant purified AK2 can efficiently process RMP in a fashion that is linear with respect to AK2 and RMP concentration in the ranges investigated (Fig. 3c, Supplementary Fig. 7). The AK2 catalytic rate (v/[E]) of 123 ([PC1]± 10) min^−1^ at 80 *μ*M RMP is only about 10-fold lower than the natural substrate AMP (Supplementary Fig. 8) and at a level that we expect to be pharmacologically relevant. These biochemical data support the proposal that AK2 phosphorylates RMP into RDP, playing a vital role in activating remdesivir to promote its antiviral efficacy and toxicity.

To understand whether loss of either gene alters mitochondrial function upon treatment with remdesivir, we measured the production of mtROS and ATP in knockout cell lines and found no significant difference in the presence of mtROS or ATP in AK2 or *SLC29A3* knockouts of HT29 and HepG2 cell lines compared to controls upon remdesivir treatment (Supplementary Fig. 3).

Naturally occurring population variation may affect the levels or functionality of proteins involved in remdesivir metabolism and toxicity with implications on relative drug efficacy and toxicity among patients. In AK2, *ATP2B1*, and *SLC29A3*, there are common eQTL variants which significantly increase or reduce gene expression in several tissues, including liver, sigmoid, and transverse colon (GTEx v8, Supplementary Table 4). Additionally, we expect that nearly 5% of individuals carry a rare loss-of-function or missense variant in one of the five genes whose knockouts decrease remdesivir cytotoxicity, some of which would be expected to reduce protein function (gnomAD v2.1.1, Supplementary Table 4). It is as-yet unclear if variation in these genes influences the clinical toxicity risks of remdesivir, and it is hard to imagine patient genotyping as a companion diagnostic for remdesivir treatment in the short-term, but such variation may provide a useful lens to study the efficacy and side effect profile of remdesivir at the population level as additional COVID-19 patient sequencing data become available.

Overall, through a genomics-based approach, this work has improved understanding of how remdesivir is metabolized and how it induces cytotoxicity. The prodrug remdesivir is thought to be taken into cells through passive diffusion^2^ Our CRISPR screen provides evidence that the first intracellular step of remdesivir activation, ester hydrolysis, occurs primarily by the lysosomal Cathepsin A (CTSA, Fig. 3d). After a further set of chemical reactions that do not appear to have strong monogenic enzymatic drivers^2^ yield RMP, our results suggest that this drug intermediate is phosphorylated to RDP by the adenylate kinase AK2, which resides in the mitochondrial intermembrane space, and this is further converted to the active triphosphate species by nucleoside diphosphate kinases. Most cells, including those used in this work, express robust levels of several other adenylate kinases including cytoplasmic AK1, mitochondrial matrix-localized AK3 and AK4, and nuclear AK6, so it is unclear why AK2 is the primary enzymatic activator of RMP.

Once formed, remdesivir-triphosphate restricts viral replication in the cytoplasm, while biochemical studies^14^ combined with our findings of remdesivir-induced mitochondrial dysfunction suggest that cytotoxicity predominantly occurs through its inhibition of mitochondrial RNA polymerase in the mitochondrial matrix. Our results suggest that targeting the bidirectional mitochondrial nucleoside transporter *SLC29A3^37^* robustly reduces remdesivir cytotoxicity in five cell lines derived from different organs while not as severely influencing remdesivir SARS-CoV-2 viral restriction. While SLC29A3 is believed to reside on both outer and inner mitochondrial membranes^37^, our findings raise the hypothesis that SLC29A3 loss may have a larger role in preventing transport of activated remdesivir through the mitochondrial inner membrane into the mitochondrial matrix, thus mitigating toxicity, without substantially limiting flow of remdesivir into and out of the mitochondrial outer membrane, which would be expected to suppress remdesivir activation by impeding AK2 access to the remdesivir prodrug.

More work is required to gauge the therapeutic relevance of SLC29A3 as a target to reduce toxicity of nucleoside drugs such as remdesivir. There are four equilibrative nucleoside transporter (ENT) genes, all of which are expressed broadly across human tissues, yet only *SLC29A3* has an effect on remdesivir toxicity. Non-specific small molecule ENT inhibitors have been identified^51^, although gene-specific inhibitors have not been identified. Homozygous germline *SLC29A3* knockout is associated with histiocytosis (macrophage expansion) in mice and humans^50,51^, although it is unknown whether transient inhibition of SLC29A3 would have side effects. To gauge the relevance of SLC29A3 as a therapeutic target, animal virus restriction studies on *SLC29A3* deficient models should be performed, and gene-specific inhibitors must be identified and tested for safety and efficacy *in vivo*.

Once properly activated, remdesivir-triphosphate’s primary effect on cellular gene expression is a reduction of mitochondrial respiratory gene expression transcribed from the nuclear genome, accompanied by reduced mitochondrial respiration and ATP production (Fig. 3c). Because remdesivir-triphosphate inhibits mitochondrial RNA polymerase^14^, the decrease in mitochondrial function may be a direct result of such inhibition as has been suggested for HCV nucleoside analogues^13^. The decrease in mitochondrial respiratory gene expression from the nuclear genome suggests that a cellular feedback mechanism that recognizes mitochondrial dysfunction and decreases mitochondrial activity to compensate may also contribute to this phenotype. In contrast to remdesivir, HIV-restricting nucleotide analogue drugs have been found to increase mitochondrial gene expression as a stress response^46^, an intriguing difference that may owe to the fact that these other drugs interfere with mitochondrial DNA polymerase in contrast to remdesivir’s impairment of mitochondrial RNA polymerase.

While mitochondrial activity is clearly central to remdesivir toxicity, our findings suggest that its role is complex, likely because of the pleiotropic role of mitochondrial regulators. For example, the CRISPR-Cas9 screening suggests that increasing the activity of NRF2 (through *KEAP1* loss) and mTOR mitigates remdesivir toxicity. Along with their role in increasing mitochondrial activity, both of these pathways promote purine biosynthesis^52,43^, a pathway whose activation should mitigate remdesivir toxicity. This pleiotropy may explain the conflicting effects on remdesivir toxicity of interventions known to increase and decrease mitochondrial activity.

It is important to stress that *in vitro* studies have been shown to be imperfect surrogates for drug toxicity seen in patients^53^. Moreover, the clinical safety profile of remdesivir remains poorly characterized^1,54,55^, and it is unknown whether the mitochondrial toxicity profile of remdesivir is the limiting factor preventing higher dosing in patients. Nonetheless, these data emphasize the utility of CRISPR-Cas9 screening to reveal genes and pathways that are subject to gene-drug toxicity interactions and to suggest actionable points of intervention to modulate remdesivir cytotoxicity.

## Methods

### Cell culture and drug titration

All cell lines were obtained from ATCC and were cultured in: McCoy’s 5A media (Thermo Fisher) + 10% FBS (Thermo Fisher) (HCT-116, HT29); DMEM (Thermo Fisher) + 10% FBS (HepG2, PLC/PRF/5, HEK293-ACE2, Huh7); RPMI (Thermo Fisher) + 10% FBS (OV-90). Information on dilution of all small molecules can be found in Supplementary Table 1.

To ascertain 50% cytotoxic concentrations (CC_50_, 11 concentrations each of remdesivir (MedChemExpress, stock concentration 50 mM in DMSO) and HCQ (SelleckChem, stock concentration 50 mM in water) were tested each in four replicates with DMSO as a control. Cells were plated at 2-4∗ 10^4/cm2 in 96-well format, and media with drug was replaced every 2-3 days. One of two methods was used to measure viability. Alamar Blue (Thermo Fisher) was used at suggested concentrations, and fluorescence was measured 15-120 minutes after staining using a Victor X5 multiplate reader (Perkin Elmer). Alternately, cells visually determined to be in range of the CC_50_ concentration were trypsinized, stained with 0.5 ug/mL propidium iodide (PI, Biolegend), and subjected to flow cytometry using a Cytek DXP11 using a consistent flow rate. Cell concentration per well was calculated from flow cytometry data using FCS Express (De Novo Software) by counting the number of live (PI-) cells per second measured by the flow cytometer between 10-30 seconds after starting sample collection.

For mitochondrial function assays, cell lines were plated at 1.25∗ 10^ 5/cm2 and treated with 0.5-1.5 *μ*M remdesivir for HepG2 and 1-3 *μ*M remdesivir for HT29 two days prior to staining. Cells were incubated in 1:500 MitoROS™ 520 stock solution (AAT Bioquest), 10 *μ*M BioTracker ATP-Red Live Cell Dye (Millipore Sigma), and respective concentrations of remdesivir for 30 minutes at 37°C. Fluorescence intensity was monitored using a Cytek DXP11 flow cytometer with 530/30 nm filter (FITC channel) and 590/20 filter (PE channel). The geometric mean of fluorescence intensity was calculated using FCS Express (De Novo Software). Within each cell line, relative fold changes for remdesivir treatments were averaged given that similar results were observed among escalating remdesivir doses.

### RNA-seq

Cell lines were plated at 6∗10^4/cm2 one day prior to drug treatment. Remdesivir and HCQ were added for 8 or 24 hours at specified doses (Supplementary Table 1), followed by RNA harvest and purification using the Qiagen RNeasy Plus mini kit. Up to 1000 ng of purified total RNA was prepared for RNA-seq using the Lexogen QuantSeq 3’ mRNA-Seq kit and sequenced using Illumina Nextseq 500 with 75-nt reads at >1.6∗10^6 reads per sample.

RNA-seq reads were mapped using the Quantseq 3’ mRNA mapping pipeline as described by Lexogen to GRCh38.p12. Briefly, reads were first trimmed using bbduk from the bbmap suite (v38.79)^56^ trimming for low quality tails, poly-A read-through and adapter contamination using the recommended parameters. Then, reads were mapped using the STAR aligner (v2.5.2b)^57^ with the recommended modified-Encode settings. Finally, HT-seq (v0.9.1) count was used to obtain per-gene counts^58^. RNA-seq data are available in GEO (GSE154936).

Within each cell line, we conducted differential expression analysis using DESeq2 1.26.0 to identify significantly differentially expressed genes for each drug treatment with respect to the DMSO condition. Samples at different time points (8 or 24h) that received the same treatment were treated as replicates. We then performed pathway enrichment analysis using GSEA as previously described by Reimand et al^59^. Genes were ranked by computing the −log_10_ p-value multiplied by the sign of the log-transformed fold-change from the differential expression analysis. This ranking was used as input to fgsea 1.12.0 to identify pathways enriched from MSigDB 7.1’s GO gene set (C5)^60^. Redundant pathways were collapsed by performing hierarchical clustering on the presence/absence of genes in the leading edge for NES > 0 and NES < 0. We defined the representative pathway for each cluster to be the pathway in that cluster with the highest absolute NES.

To compare samples across treatment conditions and cell types, we computed correlations across samples at both the gene-level and the pathway-level. On the gene level, we computed correlation of the top 2000 most variable genes that had been transformed using DESeq2’s variance stabilizing transformation. On the pathway level, we computed correlation of the NES scores of the subset of pathways identified to be significant (adjusted p-value < 0.05) in at least one cell-type/condition in the GSEA analysis. Pathways that were not significant (adjusted p-value >= 0.05) in a given cell-type/condition were assigned an NES of 0.

### CRISPR-Cas9 screening

Genome-wide CRISPR-Cas9 screening was performed using pre-made Brunello^21^ lentivirus in the CRISPR-v2 backbone (Addgene 73179-LV). HT-29 cells were plated in two replicates at a concentration of 4∗10^4/cm^2^ in 25-cm^2^ plates with 8 ug/mL of polybrene for lentivirus infection. Lentivirus was added to cells at a multiplicity of infection (MOI) of ~0.5 as determined by a titration in order to have only one gRNA copy/cell.

Two days after transduction, cells were treated with 333 ng/mL of puromycin for 3 days. Each plate was then split to six 25-cm^2^ plates, using >2∗10^7^ cells per plate, along with collection of early time point genomic DNA. Four total plates, derived from two replicate infections, were treated with either Remdesivir (1.67 *μ*M) Hydroxychloroquine (30 *μ*M), or DMSO as a control.

Genomic DNA was isolated from cells after 14 days of drug treatment or after 10 days for DMSO controls, and 20 ug of gDNA was used to amplify gRNAs using an Illumina library preparation protocol (Supplementary Note 1). Pooled samples were sequenced using NextSeq (Illumina) at the Broad Institute Sequencing Facility, using 75-nt reads and collecting greater than 5∗10^6 reads per sample.

CRISPR-Cas9 screening analysis was performed using MAGeCK package v0.5.9.3^31^. MLE beta-score was calculated by using read counts from 1000 non-targeting control gRNAs for normalization, with 5 rounds of permutation. Analysis with either pre-drug bulk population or DMSO treatment as control groups was performed, which is stated for each plot (Supplementary Table 3). To compare beta-scores across different conditions, we normalized the beta-score values by using MAGeCK cell-cycle normalization method to account for possible inconsistent cell cycles in different conditions. FLuteMLE for gene ontology (pathway+GOBP)^22^ for each fraction of candidates were performed by default setting.

### Cloning of individual sgRNAs into lentiCRISPR v2 vector

In order to create individual knock-out of the genes, we first modified the lentiCRISPR v2 backbone (Addgene 52961)^61^ to contain the “FE” -modified gRNA hairpin^62^. Next, oligonucleotides (IDT) including protospacer sequences were amplified by PCR to create homology arms and cloned into lentiCRISPR v2 FE Puro backbone through NEBuilder HiFi DNA assembly (NEB). Protospacer sequences used are listed below:

sgAK2, 5’-GTGAGGCAGGCAGAAATGGT-3’;
sgSLC29A3, 5’-GGCCAGGATGACCGTCAGTG-3’;
sgKEAP1, 5’-GACAACCCCATGACCAATCAG- 3’;
sgEIF4A1, 5’-GAAGCACATCAGAAGGCATTG-3’;
sgATP2B1, 5’-GAGAGGGCTGGAATTACTGTG-3’;
sgCTSA, 5’-GTGGTGCTTTGGCTCAATGG-3’;
sgAGPS, 5’-GCAATTTGACAGCTCATGTAG-3’;
sgFAR1, 5’-GACAGACACCACAAGAGCGAG-3’;
sgTMEM189, 5’-GCCAACACCGAGTATGACGAG-3’;
sgVPS29, 5’-GATCAAATTGCCTCTGCAACA-3’;
sgPITPNB, 5’-GCTTACTTGTTCTACAGTAG-3’;
sgATP9A, 5’-GAATGCCGATTAACTTACCAG-3’;
sgSQLE, 5’-GAAAACAATCAAGTGCAGAG-3’;
sgCtrl, 5’-GTAGCCCAGGTGTGCAGGCT-3’

### AK2 kinase activity assay

Recombinant human AK2 (Novus Biologicals) activity on remdesivir monophosphate (RMP, MedChemExpress) and adenosine monophosphate (AMP, Cayman Chemical) was measured spectrophotometrically through a pyruvate kinase/lactate dehydrogenase coupled assay that monitors NADH oxidation^50,63,64^. Kinase assays were carried out in quartz cuvettes (Hellma) in 100 *μ*L reaction mixtures containing 20mM HEPES (pH 7.3), 100 mM KCl, 10mM MgCl_2_, 2mM dithiothreitol, 1 mM ATP, 100 mM NADH, 0.5 mM phosphoenolpyruvate, 2 units of pyruvate kinase/lactate dehydrogenase mixture (Millipore Sigma), and varying concentrations of AK2 (0-40 nM). Reaction mixtures were incubated at 30° C for 2 minutes before initiation upon addition of substrate (RMP or AMP, 0-80 *μ*M). Then, individual reactions were monitored in the spectrophotometer (Beckman DU640) for 10 minutes, during which the absorbance at 340 nm was recorded every 10 seconds to track NADH consumption. For each condition, two duplicate reactions were conducted (n=2). The extinction coefficient of NADH (6.22 mM^−1^cm^−1^) was then used to calculate changes in NADH concentration, and to determine the rate of RMP or AMP consumption. We assumed that two molar equivalents of ADP were formed for each turnover of AMP and one molar equivalent of ADP was formed for each turnover of RMP. Rates were calculated from the slopes by linear regression taking data in the linear range (from the first 2 minutes) of the time course and expressed as v or v/[E]. Rate measurements in the manuscript are shown +/- standard error.

### Cultivation of SARS-CoV-2

SARS-CoV-2 (USA_WA1/2020 strain) was provided by the University of Texas Medical Branch Arbovirus Reference Collection (Galveston, TX, USA) and cultivated on VeroE6 cells (ATCC). Culture supernatants were collected three days post infection and clarified by centrifugation. Titer was calculated by serially diluting virus on VeroE6 cells, then performing a RNA FISH-based focus forming unit assay for virus infection by detection of virus mRNA (described below). All SARS-CoV-2 experiments were performed under biosafety level 4 conditions in the National Emerging Infectious Disease Laboratories BSL-4 suite at Boston University.

### Measurement of remdesivir restriction of SARS-CoV-2 infection

Control or gene knockout cell lines were treated with indicated concentrations of remdesivir or vehicle diluted in culture medium and incubated at 37° C for 1 hour prior to infection. Cells were infected at an MOI of 1 and incubated for 48 hours at 37° C then fixed in 10% neutral buffered formalin. Samples were assessed for infection efficiency using an RNA fluorescence *in situ* Hybridization (FISH) method^65^ adapted to be optimal for SARS-CoV-2. FISH probes were designed to target ORF3a and S subgenomic mRNAs and were conjugated to Cy5 fluorescent dye. Cell nuclei were stained with Hoechst 33342 (Invitrogen), and samples were imaged on a Cytation 1 Cell Imaging Multi-Mode Reader (BioTek, Winooski, VT). Images were evaluated using CellProfiler^66^ and infection efficiencies were calculated from the percentage of virus mRNA positive-cells in each sample. IC_50_ values were calculated from nonlinear regression analysis and fitting a dose-response curve GraphPad Prism Software. All assays were performed at least twice and with 3 or more replicates.

### Measurement of population variation and expression for remdesivir toxicity-modulating genes

Human population variation in the genes that modulate remdesivir toxicity was measured using the Genome Aggregation Database (https://gnomad.broadinstitute.org/) publicly available v2.1.1 exome sequence data (N=125,748 unrelated individuals). All non-synonymous variants with global allele frequency of less than or equal to 0.005 (0.5%) with functional consequence of missense, stop-gain, canonical splice site, or frameshift were included, as calculated by Variant Effect Predictor v85. All variant allele frequencies were aggregated for each of the five genes whose knockouts decrease remdesivir cytotoxicity.

To identify variants which affect gene expression levels in proteins involved in remdesivir metabolism and toxicity, we used data from GTEx Analysis Release V8 (dbGaP Accession phs000424.v8.p2). All genome-wide significant eQTLs in a single-tissue were included if present in liver, sigmoid, and transverse colon tissues. Maximum Normalized Effect Size (NES) is reported, which describes the magnitude and direction of gene expression change attributable to each variant.

## Acknowledgements

The authors thank Mandana Arbab, Grigoriy Losyev, Jaron Mercer, Andrew Anzalone, and David Liu for advice and technical assistance. The authors acknowledge funding from R01HG008754 (R.I.S.), R21HG010391 (R.I.S., C.A.C.), R01AI114814 (R.A.D.), P01AI120943 (R.A.D.), R01HG010372 (C.A.C.), R01CA74305 (P.A.C.), NWO, American Cancer Society (R.I.S.), American Heart Association (R.I.S.), Qatar Biomedical Research Institute (R.I.S.), the Agency for Science, Technology and Research Graduate Academy (G.Y.), the São Paulo Research Foundation-FAPESP n° 2019/13813-6 and 2017/25009-1 (L.C.Z.), and the Massachusetts Consortium on Pathogen Readiness (R.A.D.).

## Author contributions

Conceptualization, Methodology, Writing – Original Draft and Writing – Reviewing and Editing: R.I.S., M.C., E.A., M.C.H., L.L., C.J.D., H.C.B.C., L.C.Z., M.V., M.C., S.A.B., B.K., M.S., P.A.C., R.A.D. and C.A.C.; Investigation and Validation: R.I.S., M.C., N.C., E.A., M.C.H., C.J.D., H.C.B.C., L.C.Z. M.C., S.A.B., B.K., R.A.D. and R.J.; Software, Formal Analysis and Visualization: G.Y., R.I.S., M.C.H., L.L., H.C.B.C., B.K., M.V., J.D.F. and C.A.C.; Funding Acquisition and Supervision: P.A.C., R.A.D., R.I.S.

## References

1. Beigel, J. H. et al. Remdesivir for the Treatment of Covid-19 — Preliminary Report. N. Engl. J. Med. 0, null (2020).

2. Eastman, R. T. et al. Remdesivir: A Review of Its Discovery and Development Leading to Emergency Use Authorization for Treatment of COVID-19. ACS Cent. Sci. (2020) doi:10.1021/acscentsci.0c00489.

3. Warren, T. K. et al. Therapeutic efficacy of the small molecule GS-5734 against Ebola virus in rhesus monkeys. Nature 531, 381–385 (2016).

4. Agostini, M. L. et al. Coronavirus Susceptibility to the Antiviral Remdesivir (GS-5734) Is Mediated by the Viral Polymerase and the Proofreading Exoribonuclease. mBio 9, (2018).

5. de Wit, E. et al. Prophylactic and therapeutic remdesivir (GS-5734) treatment in the rhesus macaque model of MERS-CoV infection. Proc. Natl. Acad. Sci. 117, 6771–6776 (2020).

6. Wang, M. et al. Remdesivir and chloroquine effectively inhibit the recently emerged novel coronavirus (2019-nCoV) in vitro. Cell Res. 30, 269–271 (2020).

7. Williamson, B. N. et al. Clinical benefit of remdesivir in rhesus macaques infected with SARS-CoV-2. Nature 1–7 (2020) doi:10.1038/s41586-020-2423-5.

8. Pruijssers, A. J. et al. Remdesivir Inhibits SARS-CoV-2 in Human Lung Cells and Chimeric SARS-CoV Expressing the SARS-CoV-2 RNA Polymerase in Mice. Cell Rep. 107940 (2020) doi:10.1016/j.celrep.2020.107940.

9. Scientists to Stop COVID-19.

10. Remdesivir for the Treatment of Covid-19 — Preliminary Report. N. Engl. J. Med. (2020) doi:10.1056/NEJMc2022236.

11. Organization, W. H. WHO guidelines for pharmacological management of pandemic (H1N1) 2009 influenza and other influenza viruses. (2009).

12. Young, M. J. Off-Target Effects of Drugs that Disrupt Human Mitochondrial DNA Maintenance. Front. Mol. Biosci. 4, (2017).

13. Feng, J. Y. et al. Role of Mitochondrial RNA Polymerase in the Toxicity of Nucleotide Inhibitors of Hepatitis C Virus. Antimicrob. Agents Chemother. 60, 806–817 (2016).

14. Tchesnokov, E. P., Feng, J. Y., Porter, D. P. & Götte, M. Mechanism of Inhibition of Ebola Virus RNA-Dependent RNA Polymerase by Remdesivir. Viruses 11, (2019).

15. Moffat, J. G., Vincent, F., Lee, J. A., Eder, J. & Prunotto, M. Opportunities and challenges in phenotypic drug discovery: an industry perspective. Nat. Rev. Drug Discov. 16, 531–543 (2017).

16. Subramanian, A. et al. A Next Generation Connectivity Map: L1000 Platform and the First 1,000,000 Profiles. Cell 171, 1437–1452.e17 (2017).

17. Shalem, O., Sanjana, N. E. & Zhang, F. High-throughput functional genomics using CRISPR–Cas9. Nat. Rev. Genet. 16, 299–311 (2015).

18. Yao, X. et al. In Vitro Antiviral Activity and Projection of Optimized Dosing Design of Hydroxychloroquine for the Treatment of Severe Acute Respiratory Syndrome Coronavirus 2 (SARS-CoV-2). Clin. Infect. Dis. Off. Publ. Infect. Dis. Soc. Am. (2020) doi:10.1093/cid/ciaa237.

19. Boya, P. et al. Mitochondrial membrane permeabilization is a critical step of lysosome-initiated apoptosis induced by hydroxychloroquine. Oncogene 22, 3927–3936 (2003).

20. Subramanian, A. et al. Gene set enrichment analysis: A knowledge-based approach for interpreting genomewide expression profiles. Proc. Natl. Acad. Sci. 102, 15545–15550 (2005).

21. Wang, L. et al. A Multisite-Binding Switchable Fluorescent Probe for Monitoring Mitochondrial ATP Level Fluctuation in Live Cells. Angew. Chem. Int. Ed Engl. 55, 1773–1776 (2016).

22. Iannetti, E. F. et al. Live-Imaging Readouts and Cell Models for Phenotypic Profiling of Mitochondrial Function. Front. Genet. 10, (2019).

23. Wortel, I. M. N., van der Meer, L. T., Kilberg, M. S. & van Leeuwen, F. N. Surviving Stress: Modulation of ATF4-Mediated Stress Responses in Normal and Malignant Cells. Trends Endocrinol. Metab. TEM 28, 794–806 (2017).

24. Park, Y., Reyna-Neyra, A., Philippe, L. & Thoreen, C. C. mTORC1 Balances Cellular Amino Acid Supply with Demand for Protein Synthesis through Post-transcriptional Control of ATF4. Cell Rep. 19, 1083–1090 (2017).

25. Quirós, P. M. et al. Multi-omics analysis identifies ATF4 as a key regulator of the mitochondrial stress response in mammals. J. Cell Biol. 216, 2027–2045 (2017).

26. Espinoza, J. A. et al. The antimalarial drug amodiaquine stabilizes p53 through ribosome biogenesis stress, independently of its autophagy-inhibitory activity. Cell Death Differ. 27, 773–789 (2020).

27. Bell, S. M. et al. Ursodeoxycholic Acid Improves Mitochondrial Function and Redistributes Drp1 in Fibroblasts from Patients with Either Sporadic or Familial Alzheimer’s Disease. J. Mol. Biol. 430, 3942–3953 (2018).

28. McCreath, G., Scullion, M. M. F., Lowes, D. A., Webster, N. R. & Galley, H. F. Pharmacological activation of endogenous protective pathways against oxidative stress under conditions of sepsis. Br. J. Anaesth. 116, 131–139 (2016).

29. Benjamin, D. et al. Dual Inhibition of the Lactate Transporters MCT1 and MCT4 Is Synthetic Lethal with Metformin due to NAD+ Depletion in Cancer Cells. Cell Rep. 25, 3047–3058.e4 (2018).

30. Doench, J. G. et al. Optimized sgRNA design to maximize activity and minimize off-target effects of CRISPR-Cas9. Nat. Biotechnol. 34, 184–191 (2016).

31. Wang, B. et al. Integrative analysis of pooled CRISPR genetic screens using MAGeCKFlute. Nat. Protoc. 14, 756–780 (2019).

32. Li, W. et al. MAGeCK enables robust identification of essential genes from genome-scale CRISPR-Cas9 knockout screens. Genome Biol. 15, 554 (2014).

33. Mehellou, Y., Rattan, H. S. & Balzarini, J. The ProTide Prodrug Technology: From the Concept to the Clinic. J. Med. Chem. 61, 2211–2226 (2018).

34. Yamamoto, M., Kensler, T. W. & Motohashi, H. The KEAP1-NRF2 System: a Thiol-Based Sensor-Effector Apparatus for Maintaining Redox Homeostasis. Physiol. Rev. 98, 1169–1203 (2018).

35. Lagresle-Peyrou, C. et al. Human adenylate kinase 2 deficiency causes a profound hematopoietic defect associated with sensorineural deafness. Nat. Genet. 41, 106–111 (2009).

36. Pannicke, U. et al. Reticular dysgenesis (aleukocytosis) is caused by mutations in the gene encoding mitochondrial adenylate kinase 2. Nat. Genet. 41, 101–105 (2009).

37. Govindarajan, R. et al. Facilitated mitochondrial import of antiviral and anticancer nucleoside drugs by human equilibrative nucleoside transporter-3. Am. J. Physiol. - Gastrointest. Liver Physiol. 296, G910–G922 (2009).

38. Lade, J. M., To, E. E., Hendrix, C. W. & Bumpus, N. N. Discovery of Genetic Variants of the Kinases That Activate Tenofovir in a Compartment-specific Manner. EBioMedicine 2, 1145–1152 (2015).

39. Robbins, B. L., Greenhaw, J., Connelly, M. C. & Fridland, A. Metabolic pathways for activation of the antiviral agent 9-(2-phosphonylmethoxyethyl)adenine in human lymphoid cells. Antimicrob. Agents Chemother. 39, 2304–2308 (1995).

40. Sheahan, T. P. et al. An orally bioavailable broad-spectrum antiviral inhibits SARS-CoV-2 in human airway epithelial cell cultures and multiple coronaviruses in mice. Sci. Transl. Med. (2020) doi:10.1126/scitranslmed.abb5883.

41. Sticher, Z. M. et al. Analysis of the Potential for N4-Hydroxycytidine To Inhibit Mitochondrial Replication and Function. Antimicrob. Agents Chemother. 64, (2020).

42. Tsokanos, F. et al. eIF4A inactivates TORC1 in response to amino acid starvation. EMBO J. 35, 1058–1076 (2016).

43. Saxton, R. A. & Sabatini, D. M. mTOR Signaling in Growth, Metabolism, and Disease. Cell 168, 960–976 (2017).

44. Dean, J. M. & Lodhi, I. J. Structural and functional roles of ether lipids. Protein Cell 9, 196–206 (2018).

45. Smith, J. J. & Aitchison, J. D. Peroxisomes take shape. Nat. Rev. Mol. Cell Biol. 14, 803–817 (2013).

46. Venkatachalam, T. K., Samuel, P., Qazi, S. & Uckun, F. M. Protease-mediated enzymatic hydrolysis and activation of aryl phosphoramidate derivatives of stavudine. Eur. J. Med. Chem. 40, 452–466 (2005).

47. Murakami, E. et al. Mechanism of Activation of PSI-7851 and Its Diastereoisomer PSI-7977. J. Biol. Chem. 285, 34337–34347 (2010).

48. Cole, P. A., Grace, M. R., Phillips, R. S., Burn, P. & Walsh, C. T. The role of the catalytic base in the protein tyrosine kinase Csk. J. Biol. Chem. 270, 22105–22108 (1995).

49. Lee, Y. et al. Cloning and expression of human adenylate kinase 2 isozymes: differential expression of adenylate kinase 1 and 2 in human muscle tissues. J. Biochem. (Tokyo) 123, 47–54 (1998).

50. Figueroa, D. B. et al. Discovery of genetic variants of the kinases that activate tenofovir among individuals in the United States, Thailand, and South Africa: HPTN067. PLOS ONE 13, e0195764 (2018).

51. Boswell-Casteel, R. C. & Hays, F. A. Equilibrative Nucleoside Transporters – A Review. Nucleosides Nucleotides Nucleic Acids 36, 7–30 (2017).

52. Mitsuishi, Y. et al. Nrf2 redirects glucose and glutamine into anabolic pathways in metabolic reprogramming. Cancer Cell 22, 66–79 (2012).

53. Soldatow, V. Y., LeCluyse, E. L., Griffith, L. G. & Rusyn, I. In vitro models for liver toxicity testing. Toxicol. Res. 2, 23–39 (2013).

54. Wang, Y. et al. Remdesivir in adults with severe COVID-19: a randomised, double-blind, placebo-controlled, multicentre trial. The Lancet S0140673620310229 (2020) doi:10.1016/S0140-6736(20)31022-9.

55. Mulangu, S. et al. A Randomized, Controlled Trial of Ebola Virus Disease Therapeutics. N. Engl. J. Med. 381, 2293–2303 (2019).

56. Bushnell, B., Rood, J. & Singer, E. BBMerge – Accurate paired shotgun read merging via overlap. PLOS ONE 12, e0185056 (2017).

57. Dobin, A. et al. STAR: ultrafast universal RNA-seq aligner. Bioinforma. Oxf. Engl. 29, 15–21 (2013).

58. Anders, S., Pyl, P. T. & Huber, W. HTSeq–a Python framework to work with high-throughput sequencing data. Bioinforma. Oxf. Engl. 31, 166–169 (2015).

59. Reimand, J. et al. Pathway enrichment analysis and visualization of omics data using g:Profiler, GSEA, Cytoscape and EnrichmentMap. Nat. Protoc. 14, 482–517 (2019).

60. The Gene Ontology Consortium. The Gene Ontology Resource: 20 years and still GOing strong. Nucleic Acids Res. 47, D330–D338 (2019).

61. Sanjana, N. E., Shalem, O. & Zhang, F. Improved vectors and genome-wide libraries for CRISPR screening. Nat. Methods 11, 783–784 (2014).

62. Chen, B. et al. Dynamic imaging of genomic loci in living human cells by an optimized CRISPR/Cas system. Cell 155, 1479–91 (2013).

63. Cole, P. A., Grace, M. R., Phillips, R. S., Burn, P. & Walsh, C. T. The Role of the Catalytic Base in the Protein Tyrosine Kinase Csk. J. Biol. Chem. 270, 22105–22108 (1995).

64. Lee, Y. et al. Cloning and Expression of Human Adenylate Kinase 2 Isozymes: Differential Expression of Adenylate Kinase1 and 2 in Human Muscle Tissues. J. Biochem. (Tokyo) 123, 47–54 (1998).

65. Tsanov, N. et al. smiFISH and FISH-quant - a flexible single RNA detection approach with super-resolution capability. Nucleic Acids Res. 44, e165 (2016).

66. Kamentsky, L. et al. Improved structure, function and compatibility for CellProfiler: modular high-throughput image analysis software. Bioinforma. Oxf. Engl. 27, 1179–1180 (2011).

