## Supplementary Tables, Figures, and File for "Elucidation of remdesivir cytotoxicity pathways through genome-wide CRISPR-Cas9 screening and transcriptomics"

DRAFT

**Supplementary table 1.** Drug and small molecule concentrations given to 4 different cell lines

| Cell line | Drug Type | Drug Dose | Time point | Cell line | Drug Type | Drug Dose | Time point |
|---|---|---|---|---|---|---|---|
| HT29 | DMSO | 1:2,500 | 8 hours | HepG2 | DMSO | 1:5,000 | 8 hours |
| HT29 | DMSO | 1:2,500 | 24 hours | HepG2 | DMSO | 1:5,000 | 24 hours |
| HT29 | Remdesivir | 20 µM | 8 hours | HepG2 | Remdesivir | 10 µM | 8 hours |
| HT29 | Remdesivir | 20 µM | 24 hours | HepG2 | Remdesivir | 10 µM | 24 hours |
| HT29 | HCQ | 60 µM | 8 hours | HepG2 | HCQ | 20 µM | 8 hours |
| HT29 | HCQ | 60 µM | 24 hours | HepG2 | HCQ | 20 µM | 24 hours |
| PLC/PRF/5 | DMSO | 1:5,000 | 8 hours | HCT116 | DMSO | 1:2,500 | 8 hours |
| PLC/PRF/5 | DMSO | 1:5,000 | 24 hours | HCT116 | DMSO | 1:2,500 | 24 hours |
| PLC/PRF/5 | Remdesivir | 10 µM | 8 hours | HCT116 | Remdesivir | 20 µM | 8 hours |
| PLC/PRF/5 | Remdesivir | 10 µM | 24 hours | HCT116 | Remdesivir | 20 µM | 24 hours |
| PLC/PRF/5 | HCQ | 80 µM | 8 hours | HCT116 | HCQ | 100 µM | 8 hours |
| PLC/PRF/5 | HCQ | 80 µM | 24 hours | HCT116 | HCQ | 100 µM | 24 hours |

| Compound name | Company | Product# | M.W. (g/mol) | Stock conc. (mM) | Solvent | Dose used in HT29 (µM) | Dose used in HepG2 (µM) |
|---|---|---|---|---|---|---|---|
| Remdesivir | MedChemExpress/Selleck | HY-104077/S8932 | 602.58 | 50 | DMSO | as noted | as noted |
| HCQ | Selleck | S4430 | 433.95 | 50 | H2O | as noted | as noted |
| Trigonelline | Cayman | 11904 | 173.6 | 100 | DMSO | 200 | 200 |
| DMF (dimethyl fumarate) | Cayman | 14714 | 144.1 | 50 | DMSO | 20 | NA |
| Ursodeoxycholic acid | Cayman | 15121 | 414.6 | 80 | DMSO | 60 | 60 2 |
| Metformin | Cayman | 13118 | 165.6 | 100 | PBS | 300 | NA |
| Adenosine | Cayman | 21232 | 267.24 | 100 | DMSO | 300 | 300 |
| Syrosingopine | Sigma | SML1908-5MG | 666.71 | 15 | DMSO | 3 | 3 |
| EIDD-1931 | Cayman | 9002958 | 259.2 | 50 | DMSO | as noted | NA |

DRAFT

**Supplementary table 2.** GSEA analysis on pathway-level effects on gene expression across four liver and intestinal cell lines. Provided as an Excel sheet.

DRAFT

**Supplementary table 3.** CRISPR screen raw read counts and analysis outputs by MAGeCK-MLE pipeline. Provided as an Excel sheet.

DRAFT

**Supplementary table 4.** Three genes involved in remdesivir metabolism or toxicity (*AK2*, *ATP2B1*, and *SLC29A3*) have eQTLs which significantly increase or reduce gene expression in the liver, sigmoid, or transverse colon (GTEx v8). The maximum absolute value of expression change is reported as Normalized Effect Size (NES). Some of these variants are present in high frequencies in human populations (gnomAD v3, N=71,702 jointly-called genomes). Additionally, 4.9% of individuals across gnomAD v2.1.1 (N=125,748 jointly-called exomes) carry a rare loss-of-function or missense variant in one of the five genes associated with remdesivir metabolism or toxicity. Variants with frequencies at or below 0.5% with functional consequences of missense, nonsense, frameshift, start or stop loss, and canonical splice acceptor and donor were included, as calculated using Variant Effect Predictor v85 on GENCODE v19.

| *Most significant eQTLs in colon or liver tissue* | | | | | |
|---|---|---|---|---|---|
| **Gene Symbol** | **SNP Id** | **P-Value** | **NES** | **Tissue** | **gnomAD v3 frequency** |
| *AK2* | rs35604827 | 7.2E-09 | -0.22 | Colon - Sigmoid | 0.3487 |
| *ATP2B1* | rs11105706 | 0.000028 | -0.28 | Colon - Transverse | 0.05718 |
| *SLC29A3* | rs61851313 | 3.6E-12 | 0.49 | Liver | 0.1617 |
| *eQTLs with the largest normalized effect size expression change in colon or liver tissue* | | | | | |
| **Gene Symbol** | **SNP Id** | **P-Value** | **NES** | **Tissue** | **gnomAD v3 frequency** |
| *AK2* | rs7548117 | 9.6E-09 | -0.23 | Colon - Sigmoid | 0.2997 |
| *ATP2B1* | rs11105706 | 0.000028 | -0.28 | Colon - Transverse | 0.05718 |
| *SLC29A3* | rs61851313 | 3.6E-12 | 0.49 | Liver | 0.1617 |

| **Gene symbol** | **Aggregate population variation** |
|---|---|
| *AK2* | 0.01575 |
| *SLC29A3* | 0.01445 |
| *ATP2B1* | 0.00500 |
| *KEAP1* | 0.00601 |
| *CTSA* | 0.00812 |
| **Total** | **0.04936** |

DRAFT

### Supplementary Figures

DRAFT

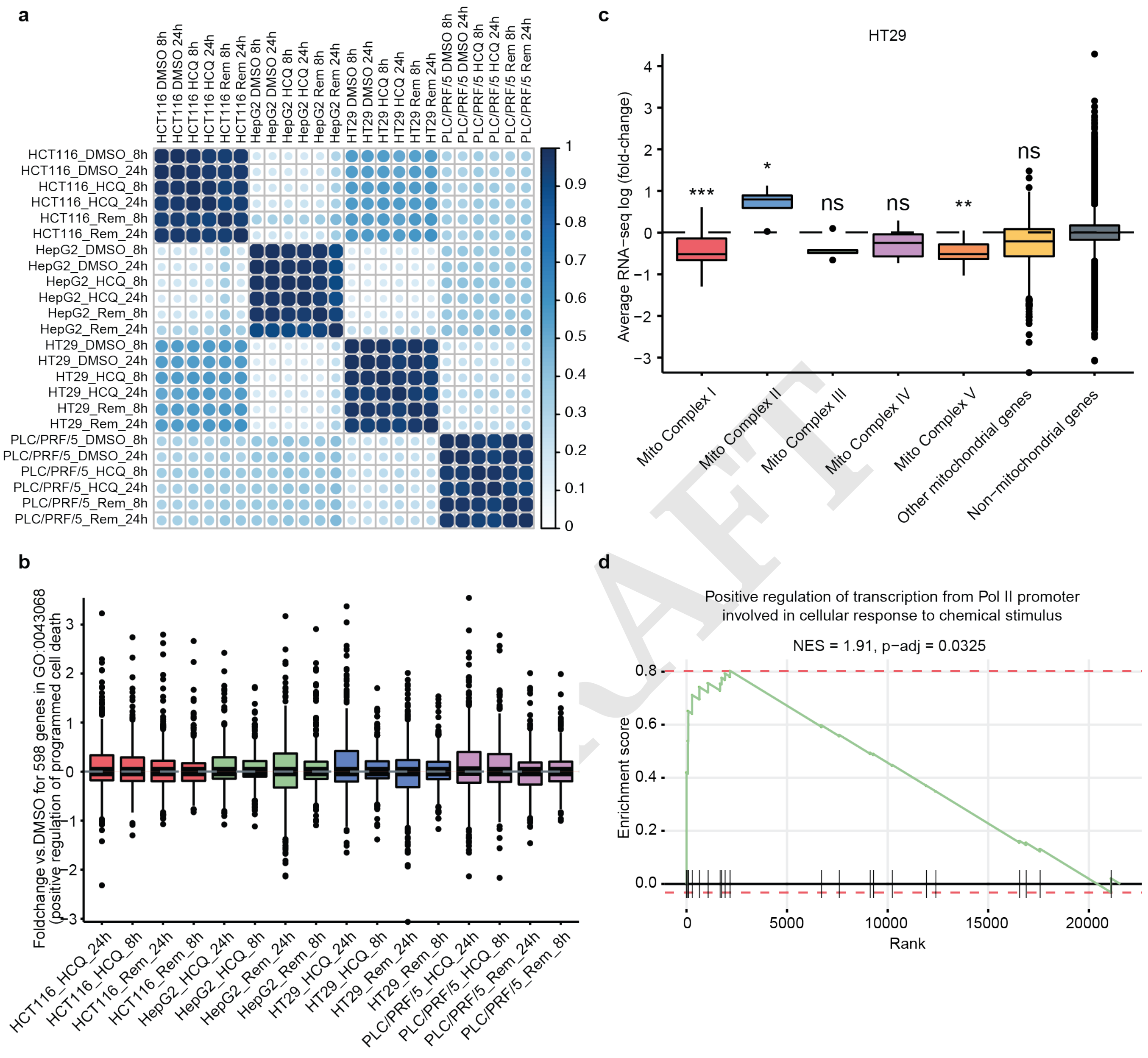


**Figure S1. GSEA reveals consistent drug-specific effects on gene expression across four liver and intestinal cell lines.** a) RNA-seq gene expression correlation matrix among HCT116, HepG2, HT29, and PLC/PRF/5 cell lines after treatment with remdesivir, HCQ, or DMSO (control) for 8 and 24 hours. b) Fold-change vs DMSO for 598 genes in GO:0043068 (positive regulation of programmed cell death) in HCT116, HepG2, HT29, and PLC/PRF/5 cell lines after treatment with remdesivir or HCQ for 8 and 24 hours. c) Average RNA-seq log(fold-change) of mitochondrial and non-mitochondrial genes in HT29 cells. d) GSEA plot for HT29 remdesivir-treated cells for genes in the GO term "positive regulation of transcription from Pol II promoter involved in cellular response to chemical stimulus."

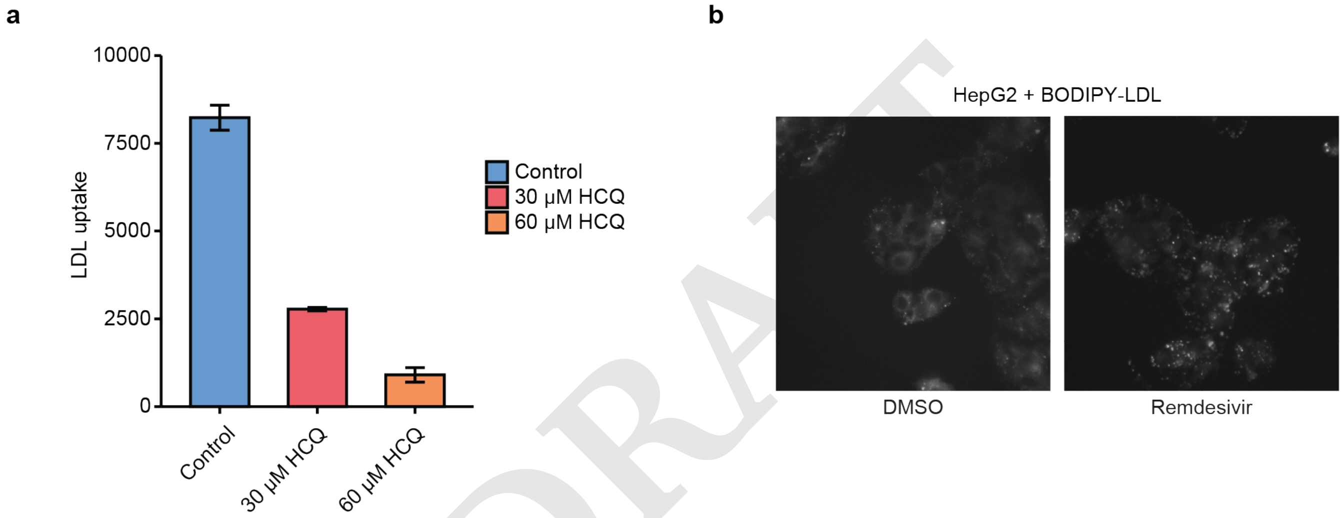


**Figure S2. Hydroxychloroquine impairs cellular LDL uptake and induces an endosomal LDL retention phenotype.** a) Uptake of BODIPY-LDL (arbitrary fluorescence units) in HepG2 cells treated with noted doses of HCQ for 24 hours. b) Fluorescent imaging of HepG2 cells after 24 hour exposure to BODIPY-LDL following treatment with DMSO or 20 $\mu$M HCQ.

DRAFT

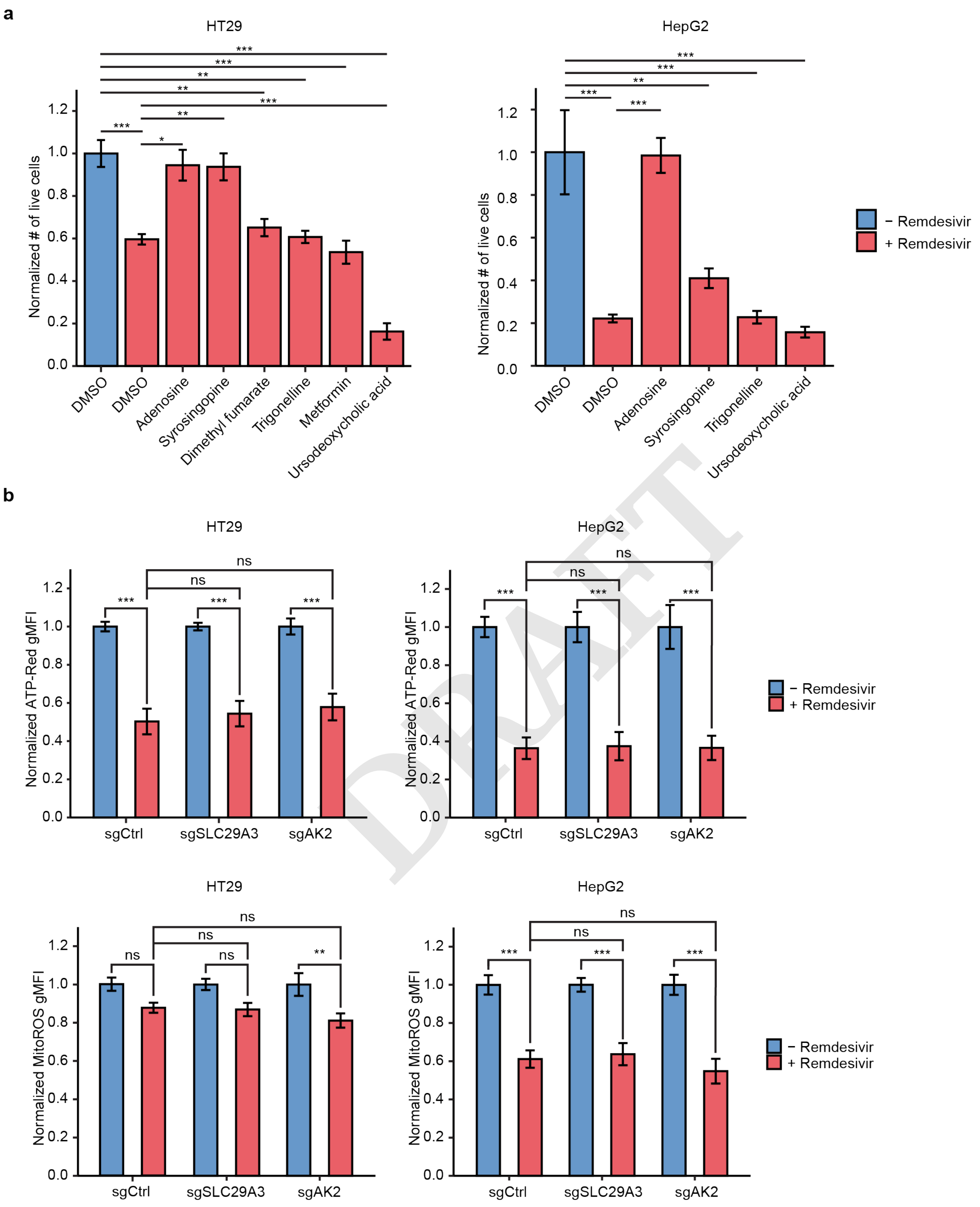


DRAFT

**Figure S3. Certain drugs that impact mitochondrial function alter remdesivir cytotoxicity.** a) Normalized number of live cells after treatment with mitochondrial drugs and with (red) or without (blue) remdesivir in HT29 and HepG2 cells. b) Normalized ATP-red and mitoROS geometric mean of fluorescence intensity (gMFI) for untreated (blue) or remdesivir-treated (red) HT29 and HepG2 control, SLC29A3, or AK2 KO cell lines.

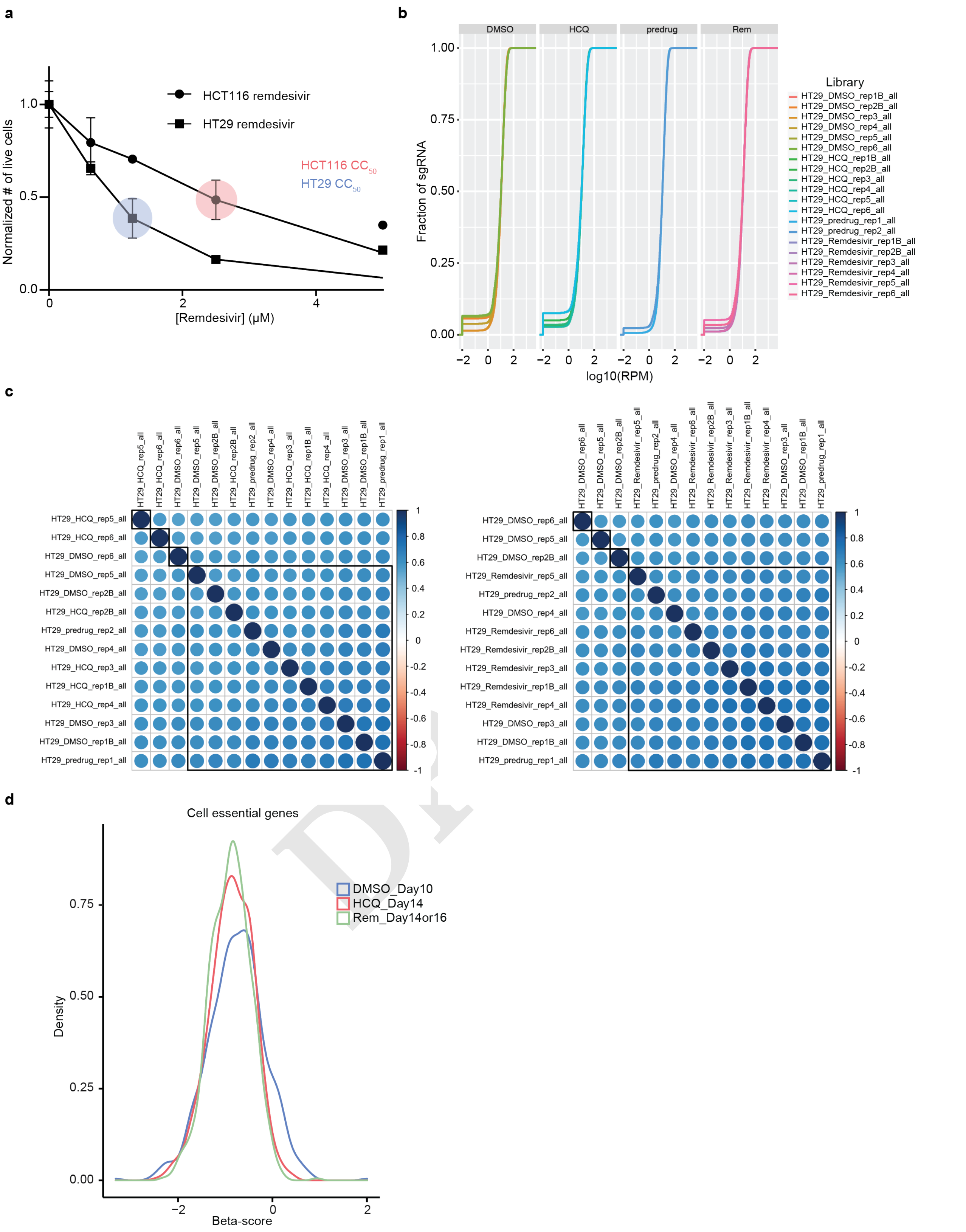


**Figure S4. CRISPR-Cas9 screening for remdesivir and hydroxychloroquine toxicity yields replicate-consistent changes in gene representation with depletion of cell essential genes.** a) Determination of the $CC_{50}$ concentrations of remdesivir and HCQ to be applied for CRISPR-Cas9 screening. b) Cumulative fraction of gRNAs from CRISPR-Cas9 screening library shown in log10-normalized read counts per million from each class of biological replicates before and after treatments with remdesivir, HCQ, or DMSO. c) Correlation matrix of gRNA readcounts in HT29 cell line before and after different treatments. d) Normalized beta-score distribution of cell essential genes after treatments with remdesivir, HCQ, or DMSO versus predrug control.

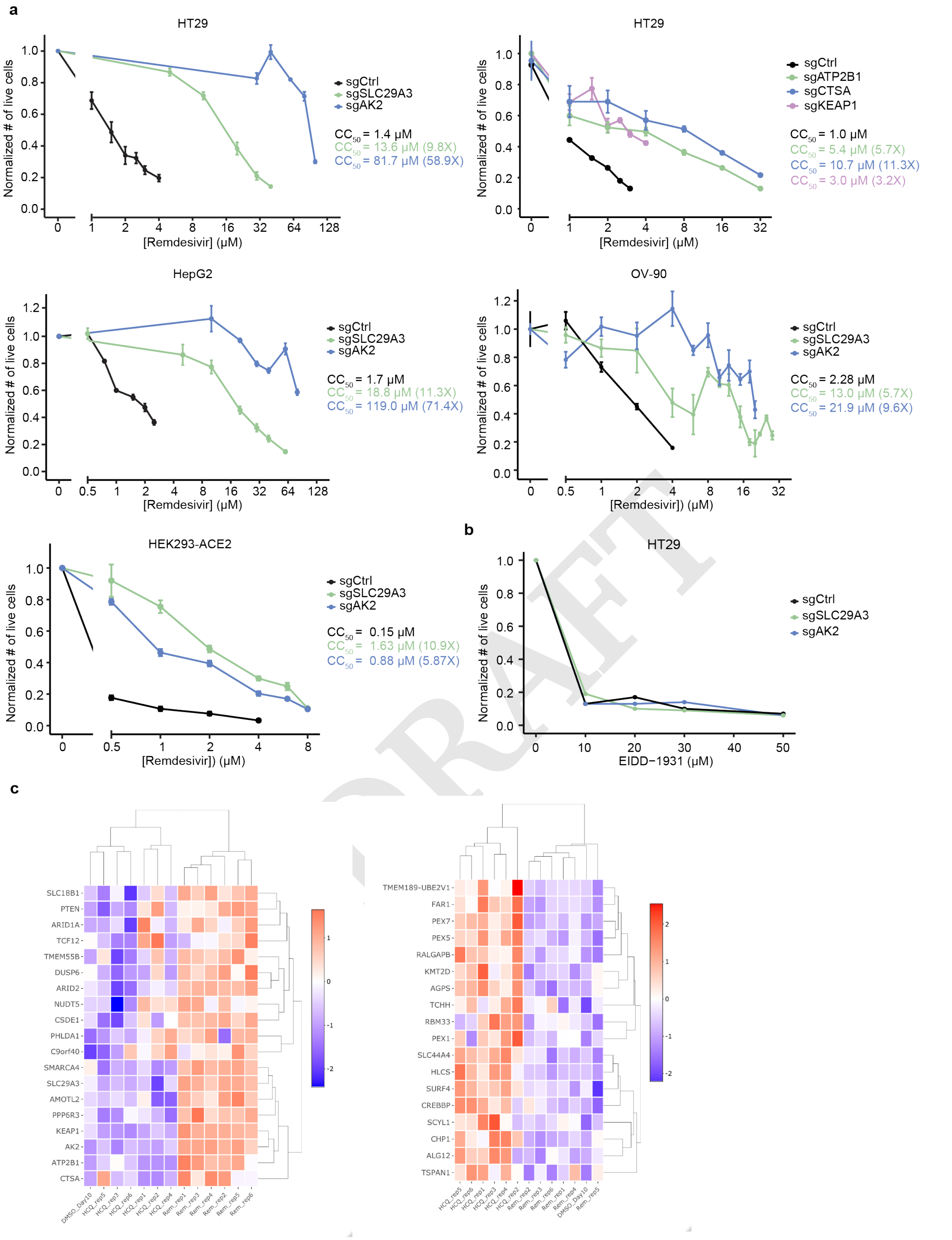


**Figure S5. CRISPR-Cas9 screening reveals genes involved in remdesivir cytotoxicity.** a) Effect of remdesivir on average normalized number of live cells and $CC_{50}$ in sgCtrl and KO cell lines in a number of cell lines. b) Normalized number of live cells after treatment with EIDD-1931 in HT29 KO cell lines. c) Heatmaps displaying MaGECK beta-scores for a set of genes that consistently mitigate remdesivir toxicity (left panel) or HCQ toxicity (right panel) in HT29 cells in each of six replicates of remdesivir treatment, six replicates of HCQ treatment, and the average of six replicates of DMSO treatment.

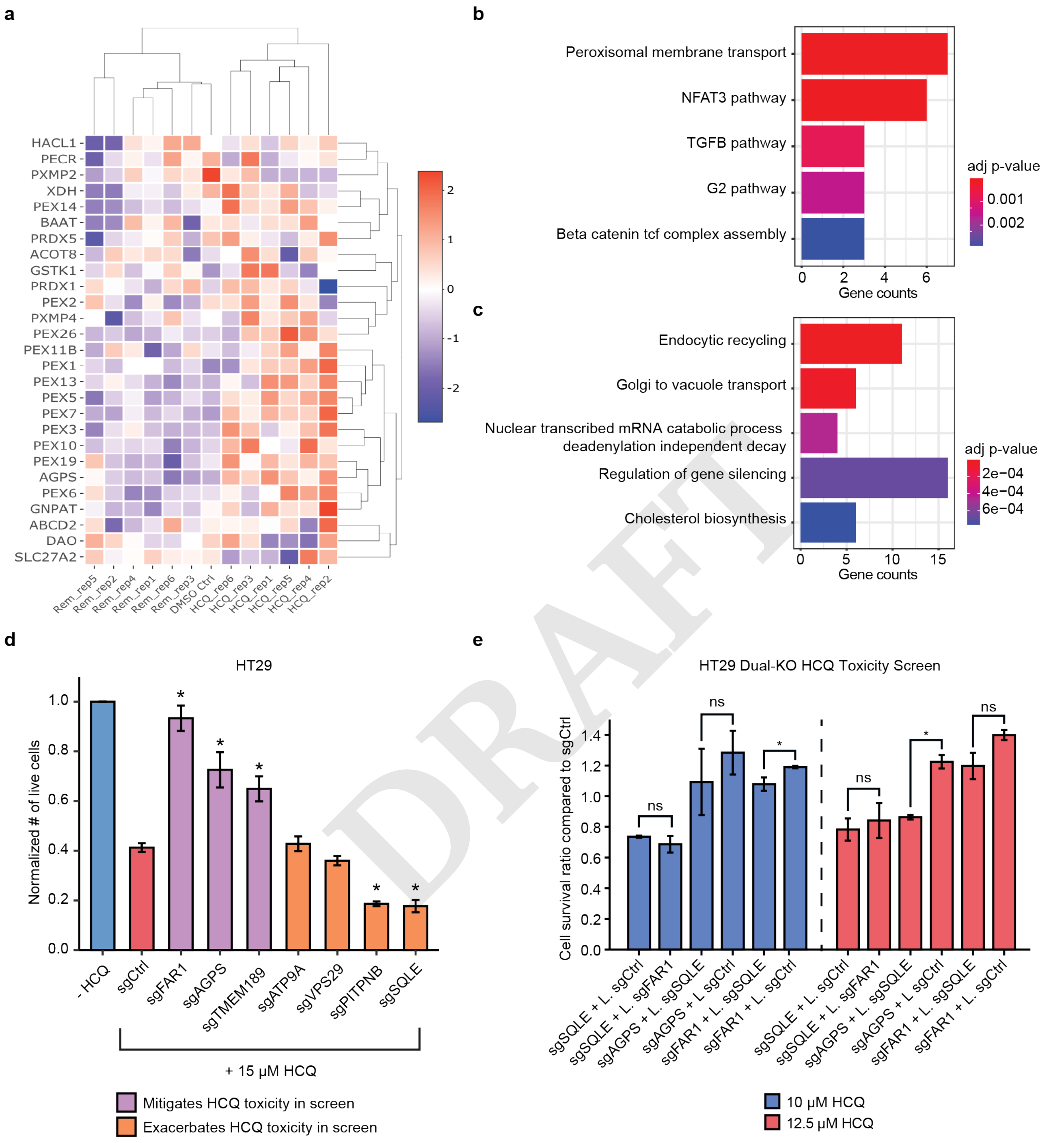


**Figure S6. CRISPR-Cas9 screening reveals that hydroxychloroquine toxicity is influenced by a coherent set of biological pathways.** a) Heatmap showing the genes involved in peroxisome KEGG pathway (hsa04146) with differentially enriched beta-score in HCQ treatments from all treatment groups. b-c) Top 5 most significant GO terms enriched (b) and depleted (c) after HCQ treatment vs. DMSO treatment. d) Effect of a $CC_{50}$ dose of HCQ on average normalized number of live HT29 cells after treatment with sgControl or KO of 7 genes whose loss was found to significantly mitigate (purple) or exacerbate (orange) HCQ toxicity in the genome-wide screen. e) Effect of two near-$CC_{50}$ doses of HCQ on average normalized number of live HT29 cells after treatment of KO cell lines (noted before +) with additional lentiviral sgControl or KO to address synthetic mitigation and exacerbation of HCQ lethality.

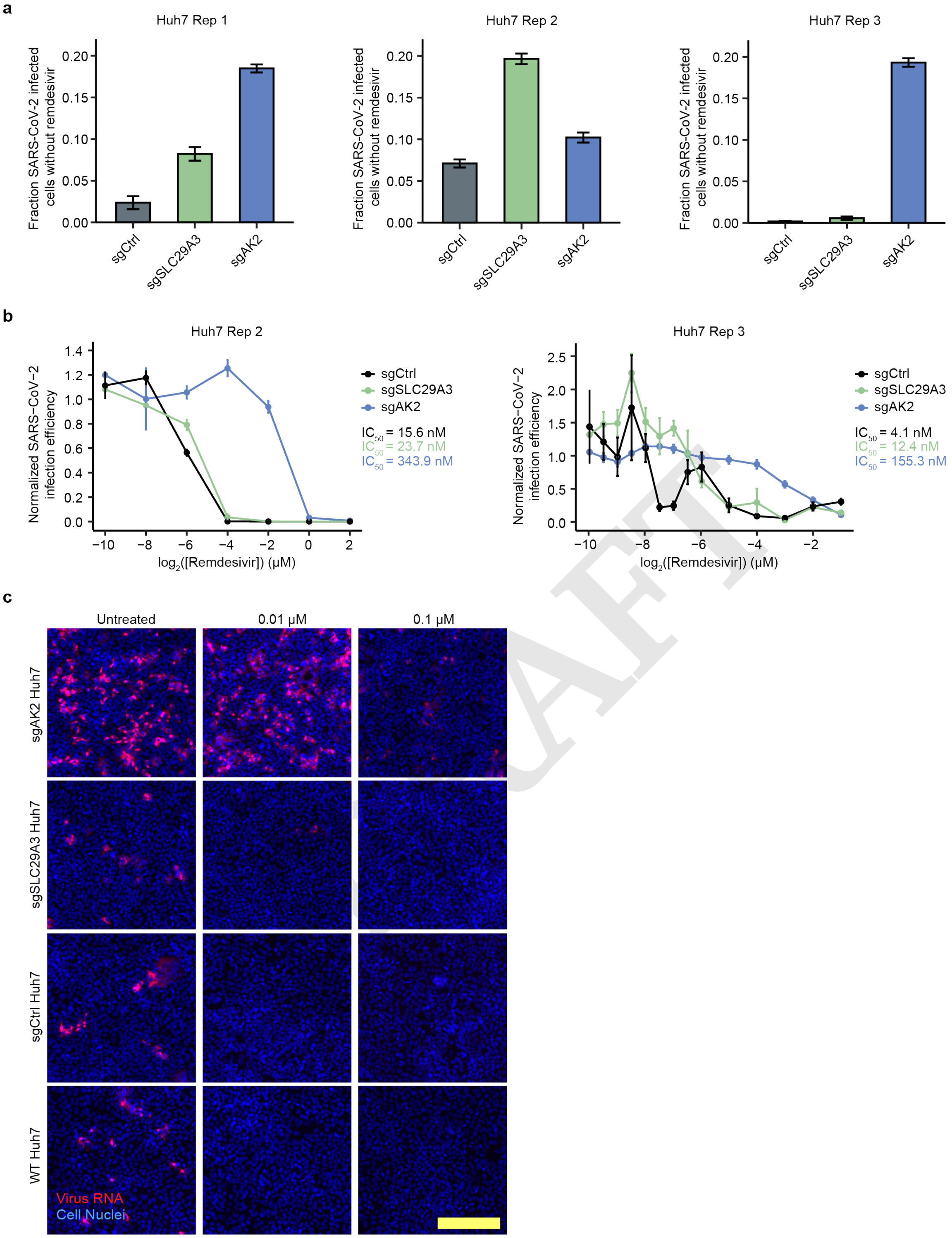


**Figure S7. Remdesivir restriction of SARS-CoV-2 infection is altered slightly in *SLC29A3* knockout and severely in *AK2* knockout cells.** a) Absolute fraction of SARS-CoV-2 infected cells in Huh7 sgCtrl and knockout cell lines in three replicate infection experiments, as measured by RNA-FISH. b) Effect of remdesivir on average normalized SARS-CoV-2 infection efficiency and $IC_{50}$ in Huh7 sgCtrl and knockout cell lines in a second and third replicate. c) Representative SARS-CoV-2 RNA-FISH images in Huh7 sgCtrl and knockout cell lines. Remdesivir concentrations are noted above the images, scale bar is 200 $\mu$m.

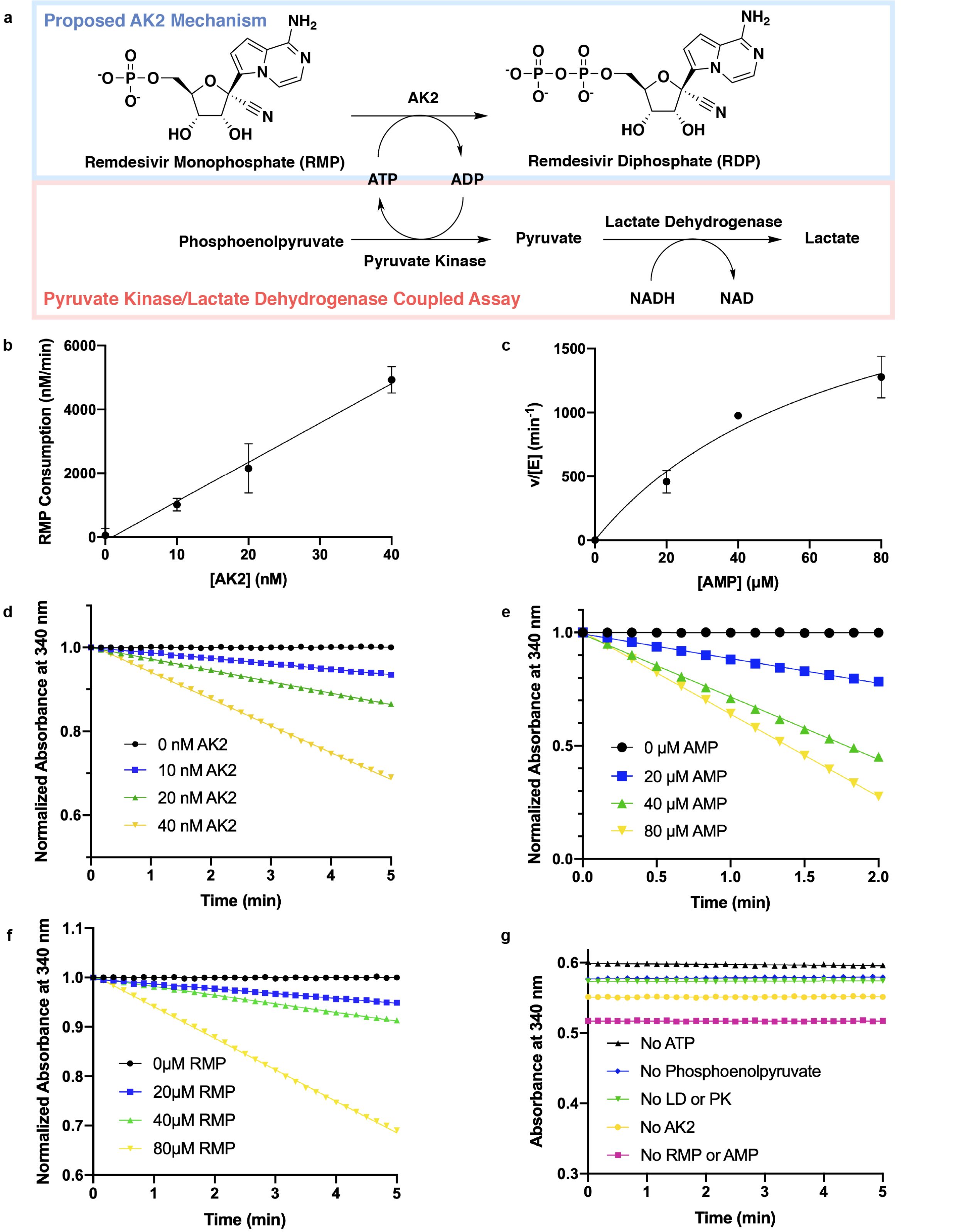


**Figure S8. Pyruvate kinase/lactate dehydrogenase coupled assay data supports phosphorylation of RMP by AK2 and allows for comparison of catalytic rates (v/[E]) between RMP and natural substrate AMP.** a) Reaction scheme of proposed AK2 mechanism and pyruvate kinase/lactate dehydrogenase coupled kinase assay. b) Average rate of RMP consumption at varying AK2 concentrations (0-40 nM). c) Catalytic rate (v/[E]) of AK2 at varying AMP concentrations (0-80 $\mu$M). At 80 $\mu$M AMP, the v/[E] is 1280 ($\pm$162) $min^{-1}$. d) Average normalized absorbance over time for a variety of AK2 concentrations (0-40 nM). All data points were scaled with respect to the starting absorbance (set as 1.0). Reactions were carried out with 80 $\mu$M RMP at 30$^\circ$C for 10 minutes. e) Average normalized absorbance over time for varying AMP concentrations (0-80 $\mu$M). All data points were scaled with respect to the starting absorbance (set as 1.0). Reactions were carried out with 10 nM AK2 at 30$^\circ$C for 10 minutes. f) Average normalized absorbance over time for varying RMP concentrations (0-80 $\mu$M). All data points were scaled with respect to the starting absorbance (set as 1.0). Reactions were carried out with 40 nM AK2 at 30$^\circ$C for 10 minutes. g) Time course of control experiments that left out components of the pyruvate kinase/lactate dehydrogenase assay including ATP, phosphoenolpyruvate, lactate dehydrogenase/pyruvate kinase mixture, AK2 enzyme, or substrate (RMP or AMP).

### Supplementary Files

**Supplementary File 1.**

*Titering of lentiviral gRNA library; LentiCRISPRv2 Brunello*

To determine the viral titer of the Human CRISPR Knockout Pooled Library; LentiCRISPRv2 Brunello (Addgene, # 73179-LV), PLC/PRF/5 (1.2x10^ 5/well), HepG2 (1.2x10^ 5/well), HCT-116 (6x10^ 4/well), and HT-29 (1.2x10^ 5/well) cells were plated into 24-well plates with their standard media including polybrene (8 ug/ml). To achieve 30-50% infection efficiency, corresponding to a multiplicity of infection (MOI) of ~0.5–1, each cell line was transduced with a varying volume (1 to 80 $\mu$ l) of viral library. The next day, the lentiviral library and polybrene including media were removed. Two days after transduction, puromycin was added to each cell media at appropriate doses (333 ng/ml for HT-29, 500 ng/ml for rest of the cell lines) for selection. 3-5 days after puromycin selection, the number of alive cells were counted and the proper virus amounts to be used in screening experiments were determined by comparing the survival rate of virus-treated and puromycin selected cells with control cell groups which were not including puromycin and virus.

*Library* cloning protocol

Genomic DNA was collected from cells after 5-16 days of treatment, 20ug of gDNA was used to amplify U6-3' to gRNA hairpin region from gDNA with different in line barcodes at PCR1 with the primers below. PCR2 was performed to add full-length Illumina sequencing adapters using internally ordered primers with equivalent sequences to NEBNext Index Primer Sets 1 and 2 (New England Biolabs). All PCRs were performed using NEBNext polymerase. Extension time for all PCR reactions was extended to 1 min per cycle to prevent skewing towards GC-rich sequences. The pooled samples were sequenced using NextSeq (Illumina) at the Broad Institute Sequencing Facility. The library prep primers were as follows:

040120_nonFEgRNA_r2seq_halftail
5' GACTGGAGTTCAGACGTGTGCTCTTCCGATCTACGGACTAGCCTTATTTAACTTGC3'

101317_U6PE1_BcX[r1]
5' ACTCTTTCCCTACACGACGCTCTTCCGATCT NNNNN GGAAAGGACGAAACACCG 3'

031317_10xr2seq_24bp_N70X_fw[r4]
5' CAAGCAGAAGACGGCATACGAGATNNNNNNNNGTGACTGGAGTTCAGACGTGTGCT 3'

110717_PE1_fulltail
5' AATGATACGGCGACCACCGAGATCTAC ACTCTTTCCCTACACGACGCTCTTC 3'

Next generation sequencing was performed using Illumina Nextseq using at least 75-nt reads and collecting >5∗10^6 reads per sample.

DRAFT
